# Local rewiring of genome - nuclear lamina interactions by transcription

**DOI:** 10.1101/685255

**Authors:** Laura Brueckner, Peiyao A Zhao, Tom van Schaik, Christ Leemans, Jiao Sima, Daniel Peric-Hupkes, David M Gilbert, Bas van Steensel

## Abstract

Transcriptionally inactive genes are often positioned at the nuclear lamina (NL), as part of large lamina-associated domains (LADs). Activation of such genes is often accompanied by repositioning towards the nuclear interior. How this process works and how it impacts flanking chromosomal regions is poorly understood. We addressed these questions by systematic manipulation of gene activity and detailed analysis of NL interactions. Activation of genes inside LADs typically causes detachment of the entire transcription unit but rarely more than 50-100 kb of flanking DNA, even when multiple neighboring genes are activated. The degree of detachment depends on the expression level and the length of the activated gene. Loss of NL interactions coincides with a switch from late to early replication timing, but the latter can involve longer stretches of DNA. These findings show how NL interactions can be shaped locally by transcription and point to a remarkable flexibility of interphase chromosomes.

## INTRODUCTION

In metazoan cell nuclei, large chromatin domains are associated with the nuclear lamina (NL) (de Leeuw et al., 2018; Gonzalez-Sandoval and Gasser, 2016; Kim et al., 2019; Lochs et al., 2019; van Steensel and Belmont, 2017). Mammalian genomes have roughly one thousand of such Lamina-associated domains (LADs), which are typically hundreds of kb or even a few Mb in size. The NL contacts of some LADs are highly consistent between cell types, while other LADs interact in cell-type specific (facultative) manners with the NL. How LAD - NL contacts are regulated is poorly understood.

Most genes inside LADs have very low transcriptional activity (Guelen et al., 2008; Leemans et al., 2019; Peric-Hupkes et al., 2010). When cells differentiate, detachment of genes from the NL often coincides with transcriptional activation, while increased NL interactions correlate with reduced transcription (Lund et al., 2013; Peric-Hupkes et al., 2010; Robson et al., 2016; Robson et al., 2017). These observations raise the interesting possibility that the NL helps to establish a repressive environment. In support of this notion, depletion of lamins can lead to derepression of specific genes (primarily in *Drosophila*) (Chen et al., 2014; Kohwi et al., 2013; Shevelyov et al., 2009); transfer of human inactive promoters from LADs to a neutral chromatin environment can lead to activation of these promoters (Leemans et al., 2019); and artificial tethering of some genes to the NL can reduce their activity (Dialynas et al., 2010; Finlan et al., 2008; Kumaran and Spector, 2008; Reddy et al., 2008).

This, however, does not rule out that the contacts of genes with the NL are the *consequence* of a lack of transcriptional activity, and vice versa, that genes detach from the NL *in response* to their activation. This was initially suggested by experiments with fluorescently tagged lacO arrays that were integrated in a locus near the NL. Tethering of the transcriptional activator peptide VP16 to these arrays caused repositioning away from the NL (Tumbar and Belmont, 2001). Similar observations were made when VP64 (a tetramer of VP16) was tethered to promoters of three distinct genes in LADs in mouse embryonic stem (mES) cells (Therizols et al., 2014). Another study found that activation of the long non-coding RNA gene *ThymoD* in mouse T cell progenitors contributed to the detachment of the neighboring gene *Bcl11b* from the NL (Isoda et al., 2017). The molecular signals that cause detachment of a locus from the NL are still poorly understood.

Analysis of NL detachment that follows forced activation of a gene has so far been limited to a handful of loci. It is thus unclear whether the observed detachment from the NL after transcription activation is universal, or limited to genes with particular features. For example, do the size of the gene and its level of transcription matter? Moreover, the previous studies of individual loci have only been based on microscopy-based assays such as Fluorescence In Situ Hybridization (FISH) or LacO tagging, and have only visualized the targeted genes themselves, but not the flanking DNA sequences. It has therefore remained unclear what the impact of these repositioning events is on the surrounding chromosomal regions. One possible scenario is that activation of a single gene inside a LAD leads to movement of the whole surrounding LAD to the nuclear interior. Alternatively, detachment could be restricted to the target gene itself or only affect some of its flanking regions. Possibly, detachment of one locus from the NL could be compensated by increased NL contacts of another locus nearby.

NL interactions have also been associated with the timing of DNA replication during S-phase. LADs typically coincide with late-replicating domains, but the overlap is not complete, particularly at the edges of LADs (Guelen et al., 2008; Peric-Hupkes et al., 2010; Pope et al., 2014). These local discrepancies are still poorly understood, but may provide important clues about the interplay between the mechanisms that establish LADs and late-replicating domains. Above-mentioned activation of genes in LADs with TALE-VP64 was accompanied by a switch from late to early replication; however, it was not analyzed how far this switch extends across the locus and how well it tracks with the changes in NL contacts (Therizols et al., 2014).

To study these issues, we took three complementary approaches. First, we used two VP16-tethering methods to activate a total of 14 different genes inside LADs, querying a variety of gene contexts. Second, we inactivated or truncated selected genes genetically to test whether they would re-attach to the NL. Third, we integrated an active transgene driven by a strong promoter into multiple LADs, and tested how this altered NL interactions of the integration sites and the flanking regions. In each instance, we used DamID to map NL interactions, enabling us to visualize the extent of NL detachment in detail along entire chromosomes. We also compared the changes in NL interactions to changes in replication timing.

## RESULTS

### Detachment of genes from the NL upon activation by TALE-VP64

We first employed a previously reported system in mouse embryonic stem (ES) cells, in which individual NL-associated genes are upregulated by means of TALE-VP64 fusion proteins that target the promoters (Therizols et al., 2014). In this system, relocation of the activated genes from the NL towards the nuclear interior was observed by FISH (Therizols et al., 2014). However, it is not known how much of the flanking DNA is involved in this detachment from the NL. We therefore repeated these experiments, but now we employed DamID mapping of Lamin B1 interactions. This method has repeatedly been shown to correspond well with FISH microscopy (Guelen et al., 2008; Harr et al., 2015; Kind et al., 2015; Peric-Hupkes et al., 2010; Robson et al., 2017), but it provides much more detailed maps of NL interactions.

We focused on two previously studied genes, *Sox6* and *Nrp1 (Therizols et al., 2014)*. In line with the reported FISH results we observed clear detachment of each gene from the NL, when activated by the corresponding TALE-VP64 construct (**Figure 1a,b middle panels**). To assess the statistical significance of these changes, we compared their magnitude to those observed throughout the remainder of the genome. Because the size of the affected region is *a priori* not known, we calculated this comparison for various window sizes between about 30 kb and 1 Mb. This resulted in domainograms (de Wit et al., 2008; Tolhuis et al., 2011) that depict the genome-wide ranking of displacement magnitudes as a function of window position as well as window size (**Figure 1a,b top panels**). We regard displacements that rank above the 95^th^ percentile or below the 5^th^ percentile (marked in shades of blue and red for decreased and increased NL interactions, respectively), and that occur locally near the targeted gene, to be highly likely due to direct effects. We note that some indirect displacements elsewhere in the genome may be expected, because the perturbations of *Sox6* and *Nrp1* may have secondary effects on gene expression.

**Figure 1.**
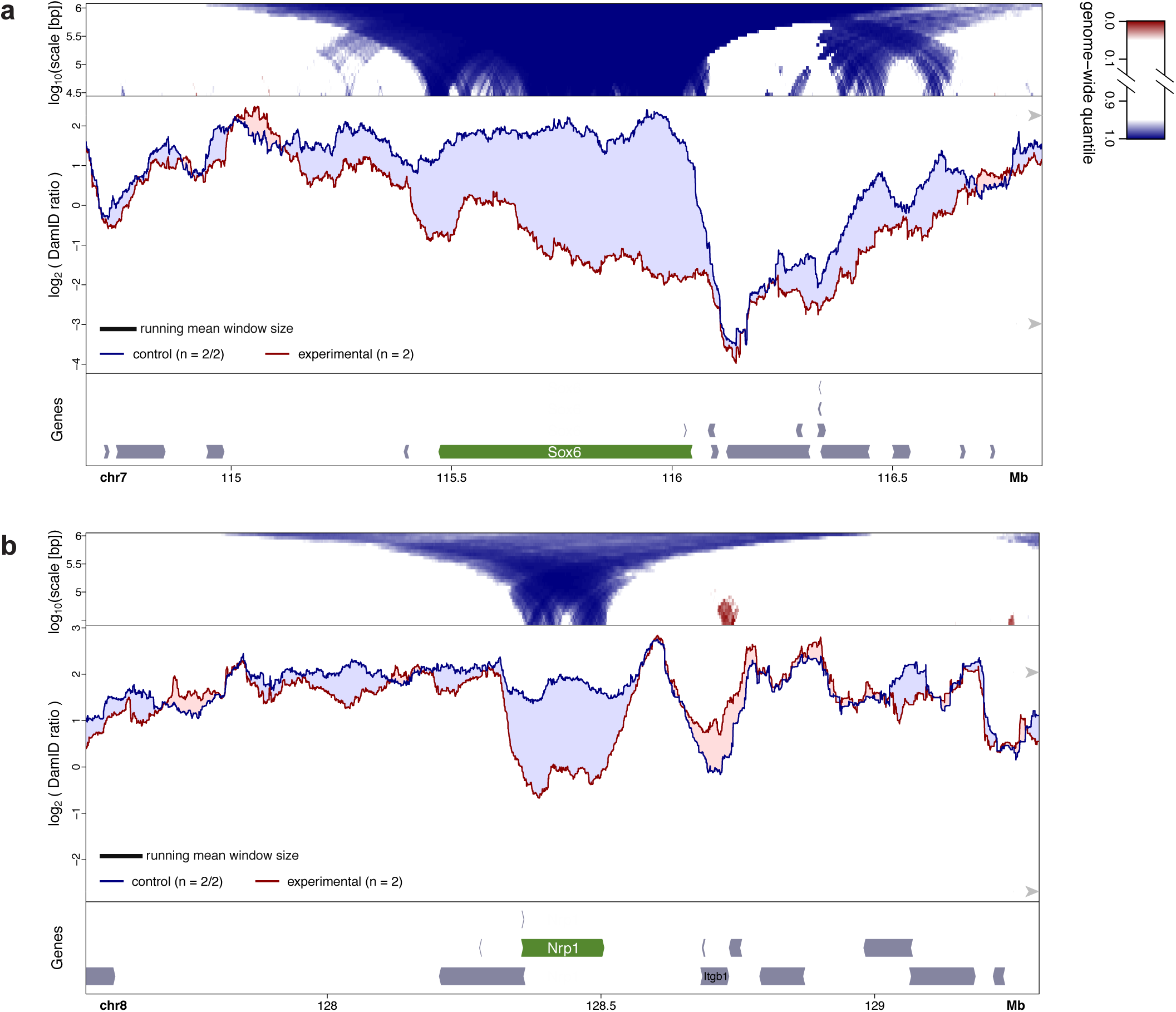
Changes in NL interactions of *Sox6* (a) and *Nrp1* (b) in mES cells after activation by TALE-VP64. *Bottom panels*: gene annotation track (mm10); the activated gene is marked in green. *Middle panels*: DamID tracks of control cells (“control”, blue line) and cells expressing TALE-VP64 (“experimental”, red line). n indicates the number of independent biological replicates that were combined. Noise was suppressed by a running mean filter of indicated window size. Shading between the lines corresponds to the color of the sample with the highest value. Arrowheads on the right-hand side mark the 5^th^ and 95^th^ percentiles of genome-wide DamID values. *Top panels*: domainograms; for every window of indicated size (vertical axis) and centered on a genomic position (horizontal axis) the pixel shade indicates the ranking of the change in DamID score (experimental minus control) in this window compared to the genome-wide changes in DamID scores across all possible windows of the same size. Blue: DamID score is highest in control samples; red: DamID score is highest in experimental samples (color key on the right of panel (a)). In **(a)** activation of *Sox6* was the experimental perturbation, activation of *Nrp1* (which is located on a different chromosome) served as control; in **(b)** activation of *Nrp1* was the experimental condition and activation of *Sox6* served as control.

The domainograms indicate that the displacements of the respective targeted genes were among the most extreme throughout the genome. For *Sox6* the NL detachment included the entire gene, but it was more pronounced near the promoter than towards the 3’ end (**Figure 1a**). Upstream of the promoter the detachment extended over ∼50-100kb, up to the LAD border. Downstream of the gene, a modest detachment was observable that tapered off over ∼300 kb. Interestingly, about 0.5 Mb upstream, across the LAD border, also some reduction in NL interactions is visible. For *Nrp1* the detachment also involved the entire gene body but did not extend much beyond it (**Figure 1b**). About 180 kb downstream of *Nrp1*, the gene *Itgb1* showed a modest increase in NL interactions. This will be discussed below. Together, these data show that activation of two genes inside LADs of mES cells results in detachment from the NL along the entire gene body, possibly with some subtler involvement of flanking regions.

### Detachment span is linked to transcript length

To extend this analysis to a larger number of genes, we switched to a more flexible gene activation system that does not require a custom-made TALE for every promoter of interest. We chose a previously established human RPE-1 cell line that stably expresses the SunCas-CRISPRa system (Tame et al., 2017; Tanenbaum et al., 2014), in which multiple copies of VP64 can be targeted to a promoter of interest by a single sgRNA. We first used this system to activate *NLGN1*, a gene of 885 kb that is located in a LAD. Transfection with a sgRNA targeting the promoter caused ∼80-fold upregulation (**Figure S1**) and resulted in clear detachment from the NL (**Figure 2a**). Relocalization primarily affected the *NLGN1* gene itself, with a mild 5’ to 3’ gradient inside the gene body and gradually decreasing along ∼100 kb of flanking DNA. We also activated the *SOX6* gene in RPE-1 cells. Again, this gene showed loss of NL interactions along its entire length, although the magnitude of the detachment was more modest (**Figure 2b**).

**Figure 2.**
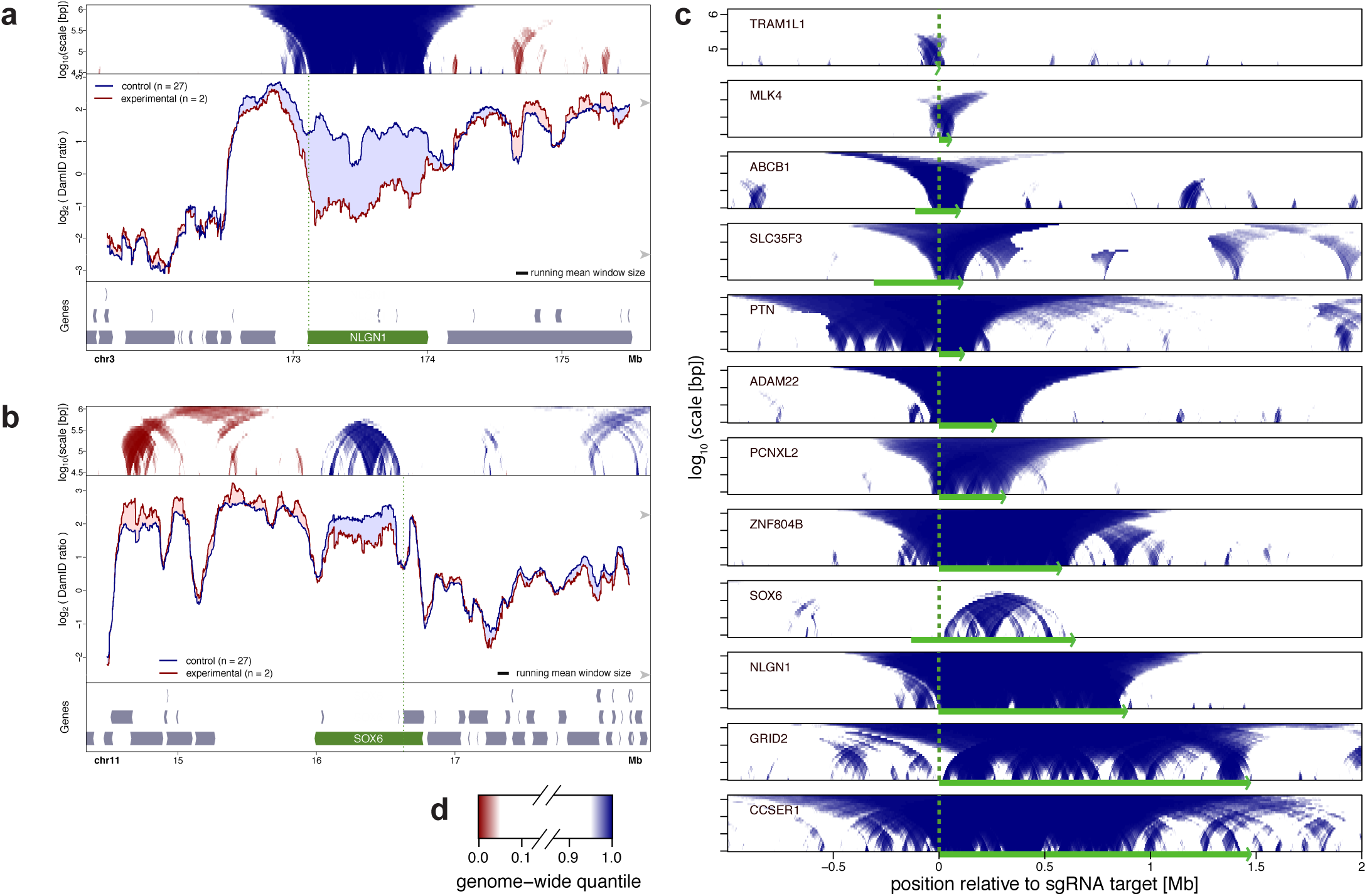
Local NL detachment caused by gene activation by CRISPRa in human RPE-1 cells. **(a-b)** Plots as in Figure 1, showing changes in Lamin B1 DamID signals upon CRISPRa activation of *NLGN1* (a) and *SOX6* (b). Control cells were treated either without sgRNA or with one of various sgRNAs targeting promoters on different chromosomes. Vertical green dotted lines mark the position of the sgRNA target sequence. **(c)** Domainograms showing regions with reduced NL interactions around 12 genes individually activated by CRISPRa. Genomic regions are aligned by the respective sgRNA target positions, and oriented so that the activated genes are all transcribed from left to right. Corresponding DamID traces are shown in **Figures 2a-b, 4-5, S3, S4**. **(d)** Color key of domainograms in **a-c**. Increases in NL interactions (red) are not shown in **(c)**.

We applied this analysis in RPE-1 cells to 12 individual genes (**Table S1**) for which activation by the SunCas system could be achieved, as determined by RT-qPCR (**Figure S1**) or RNA-seq (**Figure S2**). We chose genes of a wide variety of lengths, from ∼2kb to ∼1.5 Mb. In two cases (*ABCB1*, *SLC35F3*) we targeted a known alternative promoter located in the middle of the gene, instead of the promoter located most 5’. Strikingly, for all 12 activated genes we observed detachment of the entire region extending from the activated promoter to the 3’ end of the gene (**Figure 2c**). For most of the tested genes, detachment did not extend more than several tens of kb upstream of the activated promoter. A clear exception to this is *PTN*, which exhibited upstream detachment over nearly 0.5 Mb (see below). Likewise, for most activated genes the detachment did not extend more than 50-100 kb downstream of the 3’ end, although the precise range varied.

### Quantitative link between gene expression level and NL detachment

We wondered whether the degree of NL detachment of a gene is quantitatively linked to the transcription level. To measure gene activity accurately, we performed RNA-seq after activation of 8 of the 12 genes, and in the untreated parental cell line. Comparing RNA levels and average DamID scores across the SunCas-activated genes before and after upregulation revealed a strong negative correlation (**Figure 3**). Thus, there is a remarkably quantitative inverse link between expression levels and NL interaction frequencies.

**Figure 3.**
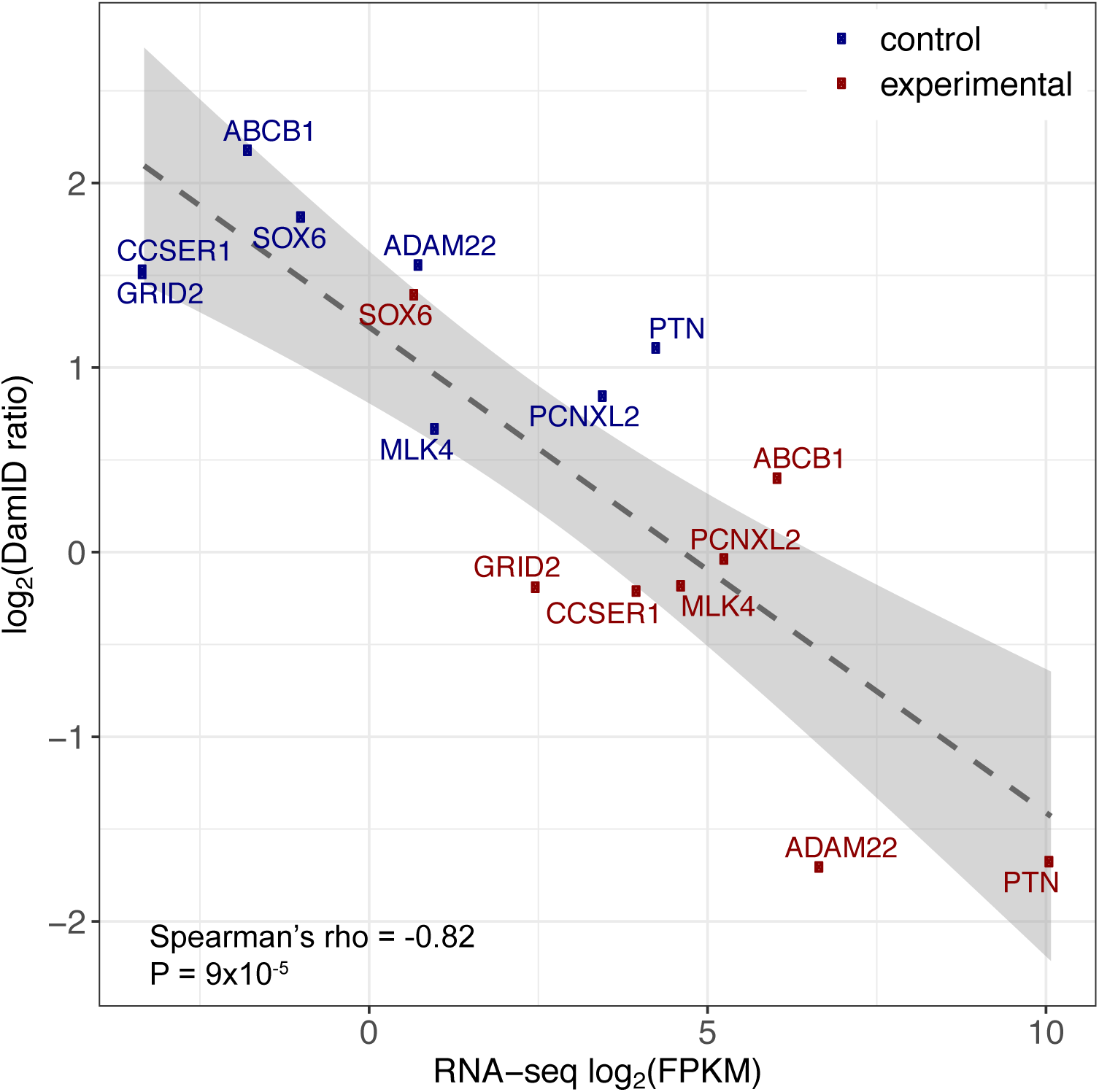
Inverse correlation between NL interaction and gene expression level. Average DamID values plotted against average expression levels of eight genes activated by CRISPRa (red; n = 2) or in control cells that were treated either without sgRNA or with one of various sgRNAs targeting promoters on different chromosomes (blue; n ranging from 19 to 27).

### Neighboring genes of targeted genes do not generally show altered expression

We also queried our RNA-seq data for neighboring genes of the activated genes. We examined genes within ∼1 Mb of our targets and generally could not detect substantial up- or downregulation (**Figure S2**). Thus, strong SunCas-induced upregulation is generally restricted to the targeted gene. Minor exceptions, of borderline statistical significance (P = 0.02, DESeq2 test) were the gene *RUNDCB3* that is partially overlapping and antisense to the activated gene *ABCB1* (**Figure S2a**), and the gene *STEAP4* nearby the activated ADAM22 gene (**Figure S2b**). However, the absolute expression levels of these co-activated genes remained much lower than their CRISPRa-targeted neighbors.

Some flanking genes co-detached from the NL together with the activated gene, but showed no detectable change in expression. The most striking example is the gene *DGKI* that flanks *PTN*. Much of this ∼0.5 Mb gene shows reduced NL interactions upon activation of *PTN*, but *DGKI* does not undergo a detectable upregulation (**Figure S4g**). We conclude that CRISPRa activation and the ensuing changes in NL contacts generally do not have substantial effects on the expression of nearby genes.

### NL detachment partially overlaps with changes in replication timing

Next, we investigated the link between changes in NL interactions and replication timing. We applied Repli-seq (Marchal et al., 2018) to visualize replication timing, revisiting five genes that exhibited NL detachment in RPE-1 cells upon activation by CRISPRa. For all five activated genes we observed a clear shift towards earlier replication. When the activated genes were relatively small (*ADAM22*, *ABCB1*, *PTN*) this shift was more or less symmetrical around the activated promoter and extended about 0.4-0.8 Mb on each side, i.e., well beyond the activated transcription units and also beyond the changes in NL interactions (**Figure 4a-c**). With longer activated genes (*CCSER1*, *GRID2,* both about 1.5 Mb long), again the shift in replication timing was strongest around the targeted promoter and extended about 0.6 Mb upstream (**Figure S3**). Downstream of these promoters the shift declined gradually towards the end of the gene, similar to the detachment from the NL. These data reveal that changes in replication timing only partially overlap with changes in NL interactions (see Discussion).

**Figure 4.**
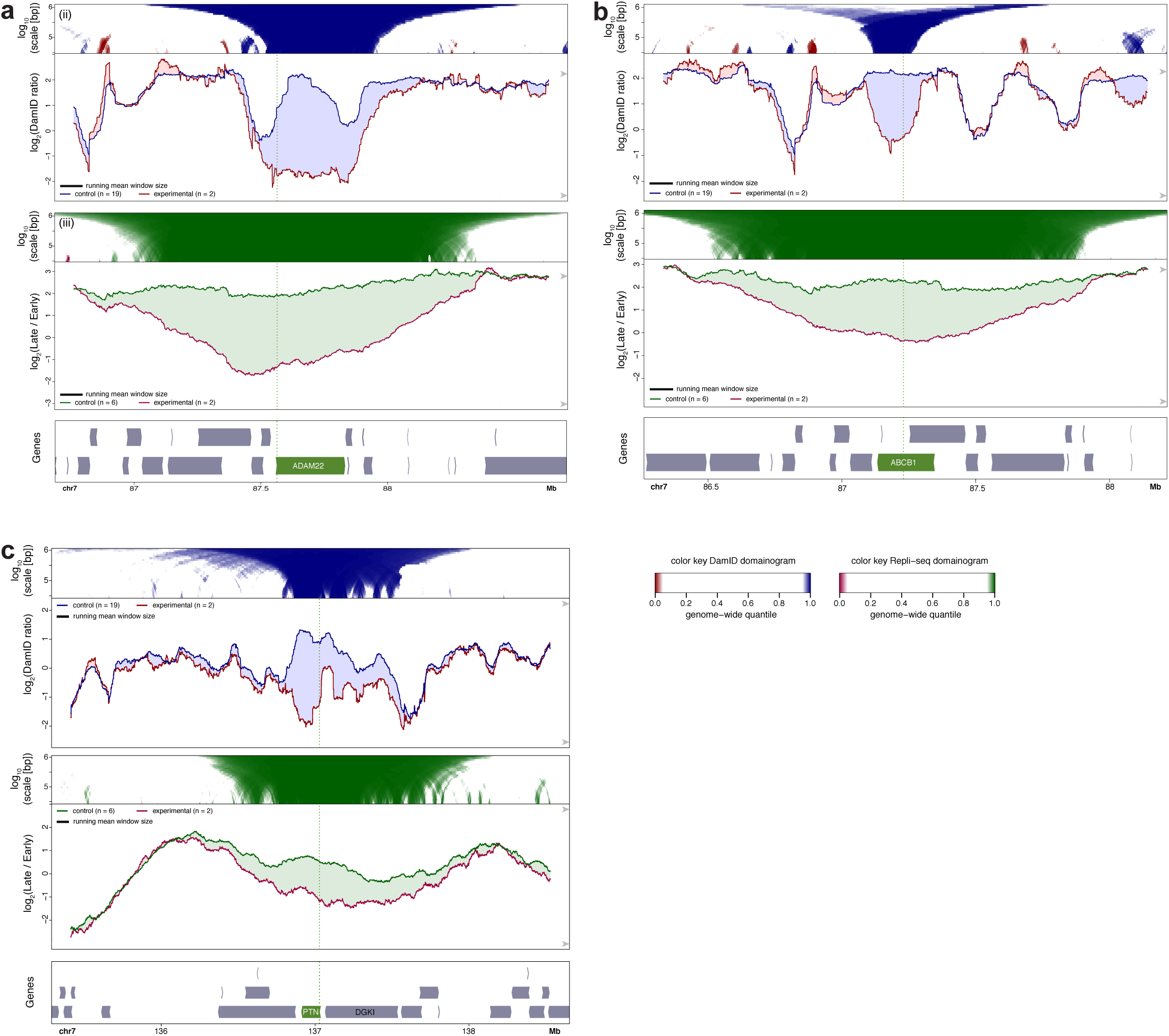
NL interactions and replication timing around activated genes. CRISPRa activation of *ADAM22* **(a)**, *ABCB1* **(b)** and *PTN* **(c)** in RPE-1 cells. Top panels show DamID data similar to **Figure 2a-b**. Middle panels show maps of replication timing at the same resolution and in the same plotting style as panels, except that different colors are used as indicated. Bottom panels show gene track, with activated gene highlighted in green.

### Possible compensatory movements

In a few instances we observed that the loss of NL interactions of the activated gene was accompanied by a gain of NL interactions of a nearby region. This was particularly notable for a region ∼0.6 Mb downstream of the activated *MLK4* gene (**Figure 5**). This region coincides approximately with gene *SLC35F3*. Although the baseline activity of *SLC35F3* is low, we found its expression to be further reduced by ∼30% (P = 0.02, DESeq2 analysis) when *MLK4* was activated (**Figure S2e**). Possibly, detachment of *MLK4* leads to compensatory movement of *SLC35F3* towards the NL, which in turn may contribute to slightly stronger repression of *SLC35F3*. Forced activation of *SLC35F3* caused its own NL detachment as expected (**Figure 2c**), but it did not alter the NL interactions of *MLK4* (**Figure S4a**). This suggests that the putative compensatory relationship is not reciprocal, but we note that this latter experiment was done only once, and should therefore be interpreted with caution.

**Figure 5.**
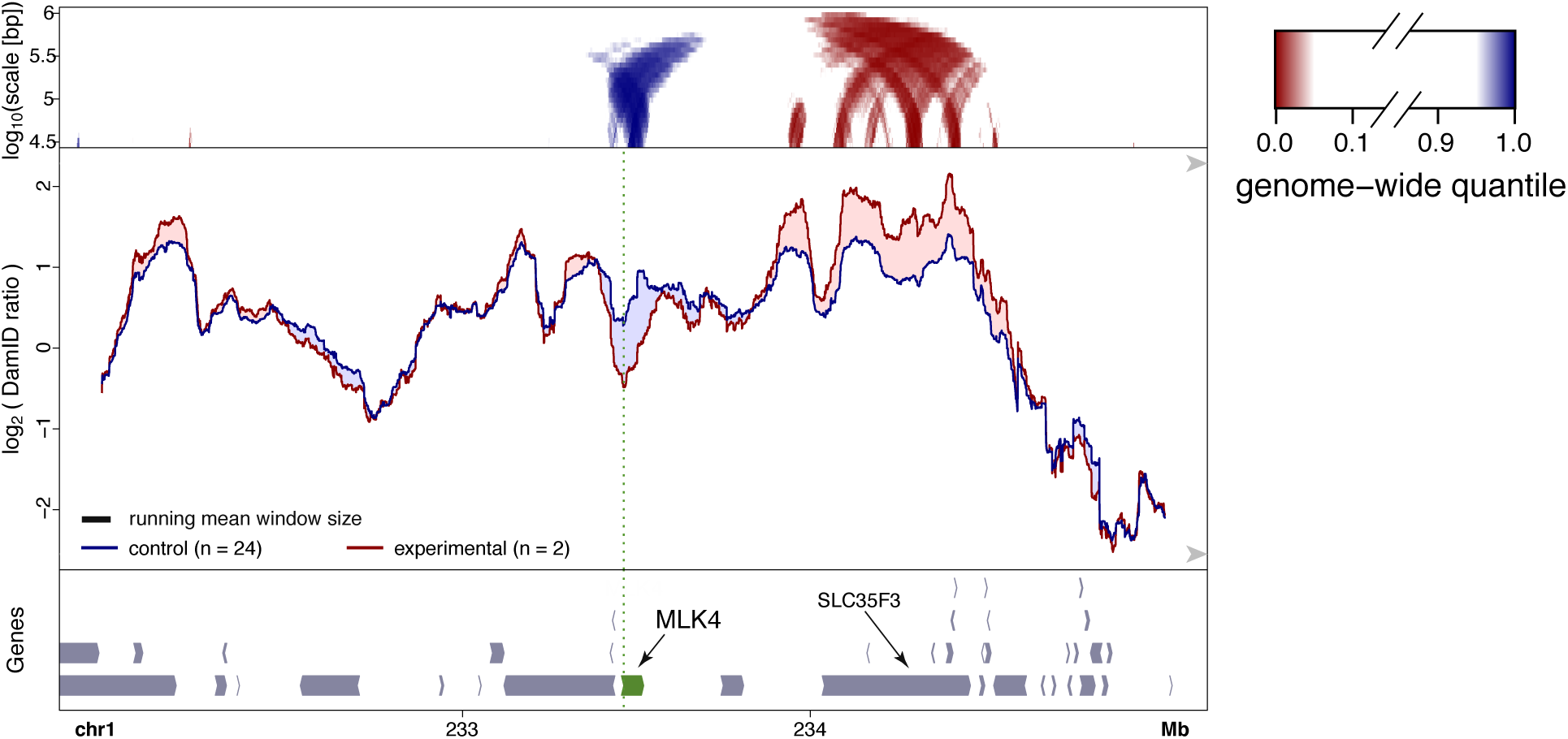
Putative compensatory movement near the activated *MLK4* gene. Changes in NL interactions after CRISPRa activation of MLK4 in RPE-1 cells. Visualization of DamID data as in **Figure 2a-b**.

We also observed moderately enhanced NL interactions of a region ∼1.3 Mb downstream of the activated *SOX6* gene (**Figure 2b**). This region coincided with the promoters of two divergent genes that were not significantly up- or downregulated (**Figure S2h**). Likewise, in case of *Nrp1* activated by TALE-VP64, the gene *Itgb1* (about 180 kb downstream of *Nrp1*) showed a modest increase in NL interactions (**Figure 1b**), but its expression was not found to be detectably altered by TALE-VP64 targeting of *Nrp1* (Therizols et al., 2014). We found also minor local increases in NL interactions within ∼2 Mb of the activated genes *NLGN1* (**Figure 2a**), *TRAM1L1* and *ZNF804* (**Figures S4b,c**). However, because of their modest magnitude we did not further investigate these movements. In summary, possible compensatory changes in NL interactions around activated genes are relatively modest, and may only anecdotally affect gene expression, at least in the cell systems we studied. These increases in NL interactions may reflect compensatory movement to fill up space at the NL vacated by the activated genes.

### No evidence for a cooperative mechanism of detachment

We next explored whether it is possible to detach a whole LAD by activating multiple genes in the domain. We first tested this for two neighboring large genes, *CCSER1* (1475 kb) and *GRID2* (1468 kb). Activation of each gene individually caused clear detachment from the NL (**Figure 6a; Figure S3a,b**). When both genes were activated simultaneously (by co-transfection of the respective sgRNAs), both genes detached, but the intervening ∼700 kb region showed no significant reduction in NL association (**Figure 6a**). Under this double-activating condition the intervening region also continued to be replicated a bit later in S-phase than the two activated genes (**Figure S3c**).

**Figure 6.**
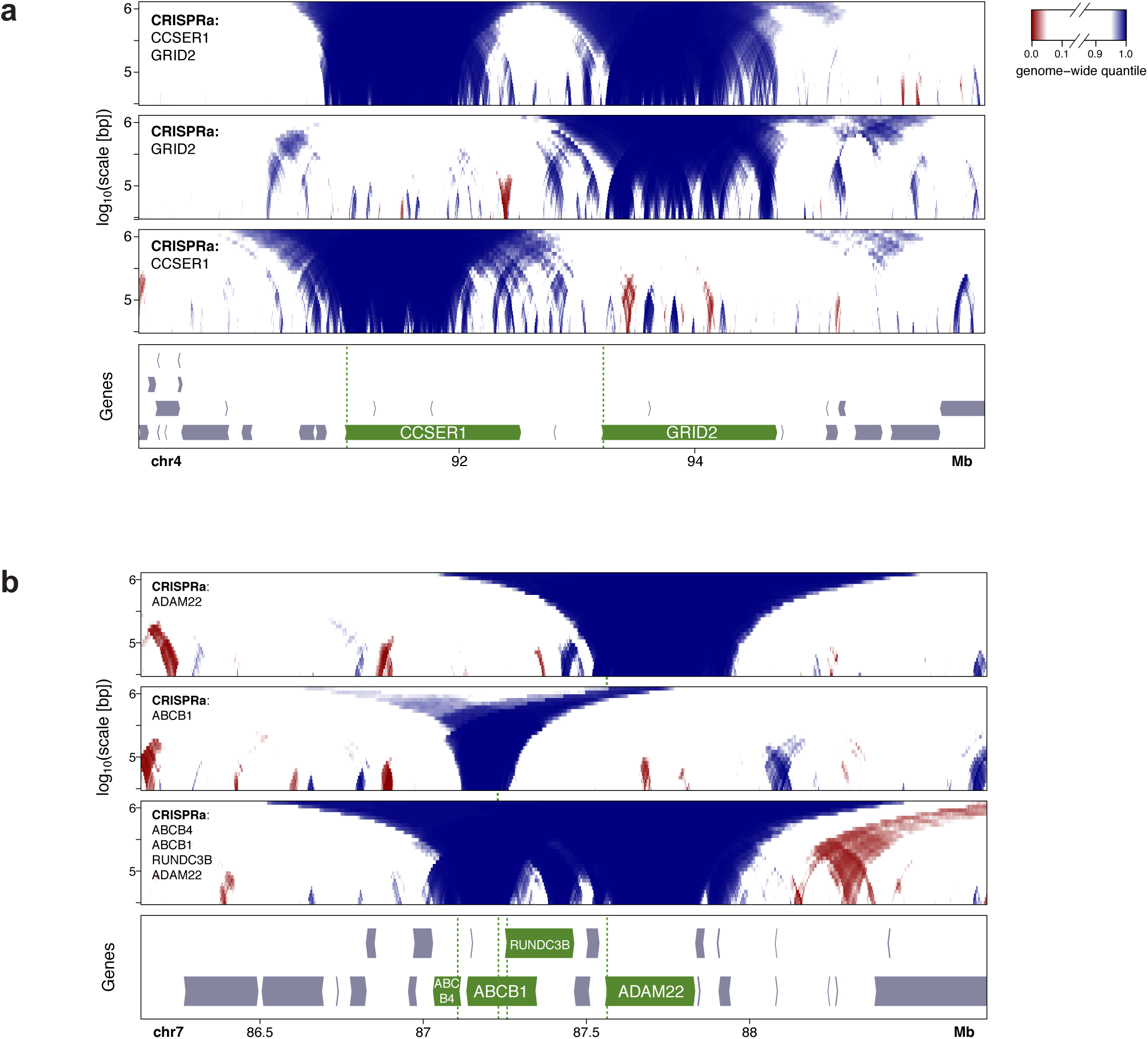
Effects of activation of multiple neighboring genes on NL interactions. **(a)** DamID domainograms after activation of *CCSER1*, *GRID2*, or both. **(b)** DamID domainograms after activation of ADAM22, ABCB1, or the genes *ABCB4*, *ABCB1*, ADAM22 and *RUNC3B* simultaneously. DamID data of activation of *ADAM22* and *ABCB1* alone are same as in **Figure 4a, b**.

We also applied CRISPRa simultaneously to the more closely spaced genes *ABCB4*, *ABCB1*, ADAM22 and *RUNC3B* by co-transfection of four sgRNAs. All four genes were induced to varying levels (**Figure S5a,b**). We compared the resulting DamID maps to those obtained after activation of ABCB1 or ADAM22 alone (**Figure 6b; Figure S5c-e**). While the single gene activations resulted in selective detachment of the respective genes, the quadruple activation caused detachment of each gene, with the degree of detachment roughly corresponding to the activity of each gene after activation (**Figure S5a, c-e**). There was no indication that the observed detachment of the four genes involved a more extensive region than a simple combination of their independent detachments. Thus, at least in the contexts tested here, detachment from the NL does not happen in a cooperative fashion.

### Inactivation of genes can promote NL interactions

Our observations so far strongly suggested that the act of transcription is a driving force that localizes genes to the nuclear interior. To test this further, we set out to block transcription by two complementary genetic strategies.

In the first strategy, we aimed to disrupt all transcription in an inter-LAD region (iLAD), to test whether this would lead to increased NL interactions of the entire region. We focused on the mouse genes *Dppa2, Dppa4, Morc1* and *Morc* (a shorter form of *Morc1* that initiates from an alternative transcription start site). In ES cells these genes are localized in an approximately 500 kb-sized iLAD. However, this region is NL-associated in mouse neural precursor cells (Peric-Hupkes et al., 2010) and therefore it has the potential to become a LAD. We used recently reported (Sima et al., 2019) F1 hybrid *Cast*/*129Sv* mES cell clones (named E2 and A6) with a heterozygous triple deletion of the promoters of *DppA2*, *Morc1* and *Morc* on the *129Sv*-derived chr16. These deletions also stop transcription of the *Dppa4* gene and therefore essentially abolish transcriptional activity in the whole iLAD (Sima et al., 2019). Owing to the high density of SNPs that differ between the *129Sv* and *Cast* genomes, we could generate allele-specific DamID maps, enabling us to compare the mutated and wild-type chromosomes.

DamID profiles of the locus revealed that *Morc1* and *Morc* on the mutated chromosome had moved towards the NL in the mutant cells, when compared to control cells carrying an unrelated mutation on a different chromosome (**Figure 7a**). This effect was not observed for the wild-type *Cast* chr16 in the same cells (**Figure 7b**). Interestingly, the region containing *Dppa2* and *Dppa4* was unaffected and clearly remained detached from the NL. To determine if ablation of the most prominent transcript would be sufficient to induce attachment, we also tested a single deletion of the *Morc* promoter (clones A12 and B11), which reduces transcription of *Morc1* by ∼2-fold and presumably ablates expression of *Morc,* but does not alter expression of *Dppa2* and *Dppa4 (Sima et al., 2019)*. In the mutated *129Sv*-derived locus this perturbation resulted in a more restricted increase in NL interactions of *Morc* while the 5’ end of *Morc1* was much less affected (**Figure 7c**). Again, in the wild-type *Cast*-derived locus only minor changes were observed between mutated and control clones (**Figure 7d**). These data show that inactivation of one or more genes in a facultative iLAD can lead *in cis* to locally increased NL interactions of the inactivated genes.

**Figure 7.**
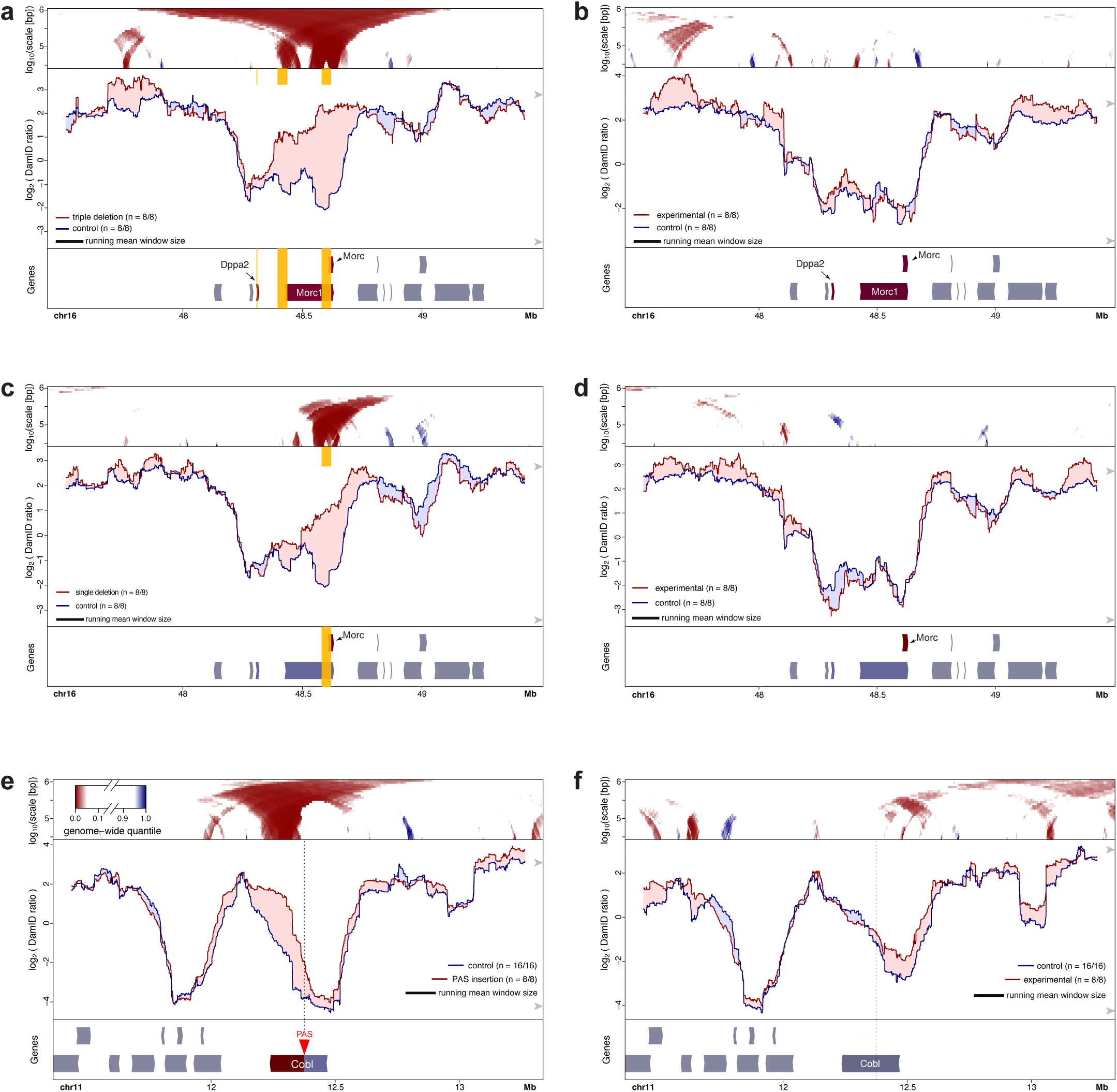
Increased NL interactions upon allele-specific transcription inactivation in F1 hybrid mES cells. **(a)** DamID profiles of the *129Sv* allele of the *Morc1* locus with deletions of the promoters of *Morc*, *Morc1* and *DppA2* (yellow boxes), and in control cells. **(b)** Same as (a), but for the (non-mutated) *Cast* alleles. **(c-d)** Same design as (a-b), but with only a single mono-allelic deletion of the *Morc* promoter (yellow box). **(e-f)** Effect of PAS insertion on NL interactions of *Cobl* gene locus. Same design as (a-b) but with insertion of a PAS (located at the green vertical dotted line) that truncates the *129Sv* allele of the *Cobl* transcription unit. *129Sv* allele is shown in (e), *Cast* allele in (f). Clones with *Morc1* locus mutations (each assayed in four independent biological replicates) served collectively as control in (e-f), and the clone with the PAS integration (eight independent biological replicates) served as control in (a-d). Visualization of DamID data in all panels is as in **Figure 1**.

In the second genetic approach, we aimed to truncate a single transcript, to test directly whether transcription elongation is required for detachment from the NL. We chose the 228 kb *Cobl* gene, which is active and locally detached from the NL in mES cells but inactive and NL-associated in neuronal precursor cells (NPCs), indicating that its detached state is facultative and linked to transcription. We created a heterozygous truncation of the *Cobl* transcription unit in F1 hybrid *Cast*/*129* mES cells by insertion of a polyadenylation sequence (PAS) in the *129* allele of *Cobl*, 89 kb downstream of the TSS. Analysis of *Cobl* allelic sequence variants in mRNA-seq data confirmed the premature termination of transcription at the *129Sv* allele (**Figure S6**). Allele-specific DamID profiles show increased NL interactions of the *129Sv* allele of *Cobl*, particularly downstream of the PAS integration (**Figure 7e**). This did not occur at the unmodified *CAST* allele, although some modest changes in NL interactions were detected in the surrounding region (**Figure 7f**). We conclude that blocking of *Cobl* transcription elongation causes local increases in NL interactions.

### Insertion of a small active gene causes moderate detachment from the NL

Finally, complementary to the activation and inactivation of genes in their native context, we tested whether insertion of a highly active transgene into a LAD is sufficient to cause local detachment from the NL. For this purpose, we designed an expression cassette consisting of a transcription unit encoding enhanced green fluorescent protein (eGFP) driven by the strong human *PGK* promoter, cloned into a PiggyBac transposable element vector (**Fig 8a**). We integrated this cassette randomly in the genome of F1 hybrid mouse ES cells by co-transfection with PiggyBac transposase. We then isolated clonal cell lines and focused on two with a large number of integrations, reasoning that by random chance several integrations would occur inside LADs. Indeed, by inverse-PCR and Tn5 mapping (see Methods) we found 26 integrations to be inserted inside LADs, out of a total of 109 in the two cell clones combined (**Fig 8b**). In comparison to the corresponding wild type alleles in the same cells, a roughly 2-fold reduction in average DamID signal was detected around the integration sites, spanning approximately 20 kb on each side (**Figure 8c**). We conclude that the integrated transcription units tend to detach the directly flanking DNA from the NL, but only partially and within a range of roughly 20 kb.

**Figure 8.**
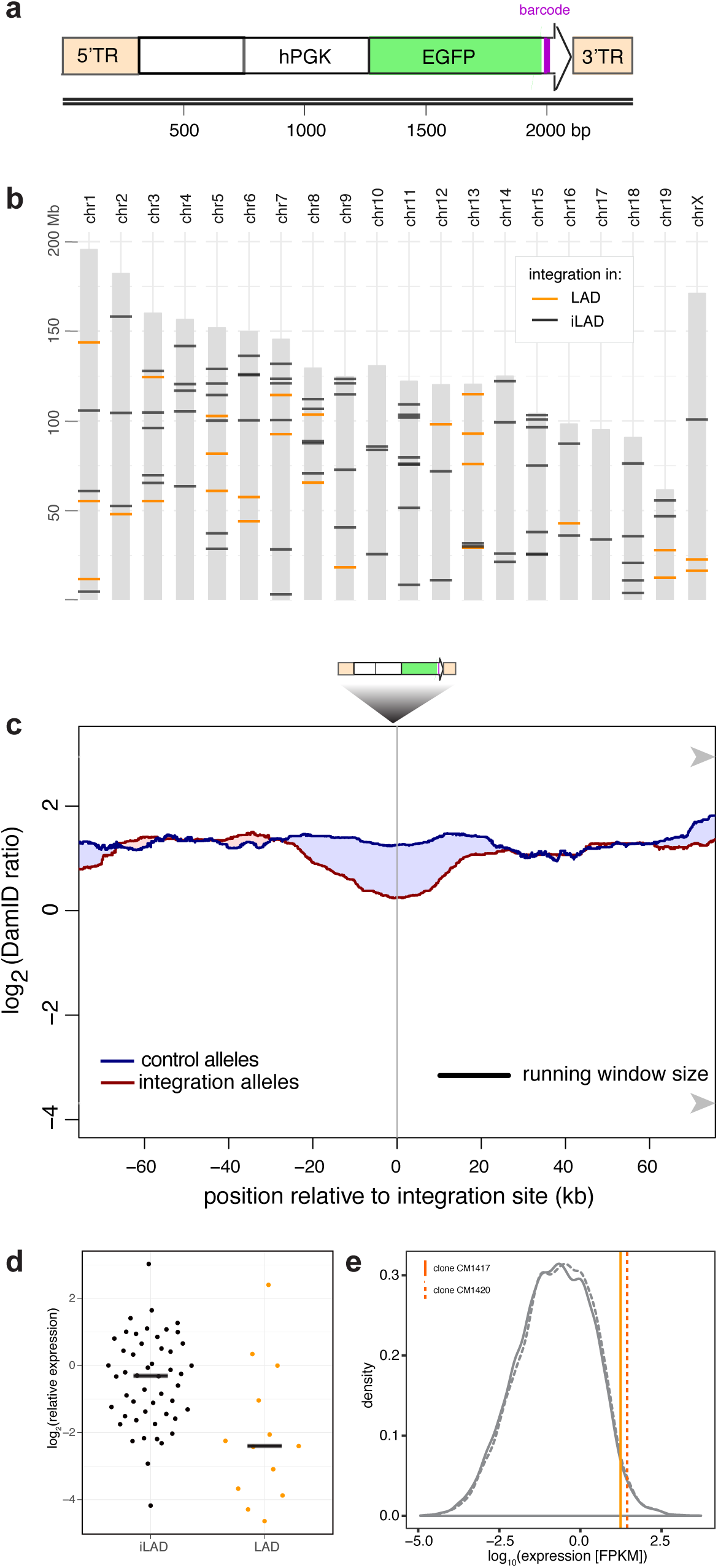
Effects of a highly active integrated transgene on NL interactions. **(a)** Design of the transgene construct, consisting of an enhanced green fluorescent protein (EGFP) transcript, marked at its 3’ end by a random barcode (purple bar) and driven by the human *PGK* promoter. The construct is flanked by the terminal repeats (5’TR and 3’TR) of the *Piggybac* transposon that are used for random integration in the genome. **(b)** Summary of the mapped locations of 109 integration sites in the genomes of two F1 hybrid mES cell clonal cell lines. LAD and iLAD integrations are shown in orange and black, respectively. **(c)** Average DamID profiles across 26 transgene integration sites inside LADs. Blue curve shows alleles without integrations, red curve shows the corresponding alleles with integrations. Shading between the lines shows which curve has the highest value. **(d)** Relative expression levels of individual barcoded transgenes in LADs and iLADs. **(e)** Estimated average expression level of integrated transgenes in LADs for the two clonal cell lines CM1417 and CM1420, compared to the distribution of expression levels of all active endogenous genes.

We considered the possibility that our expression cassette was not strong enough to cause more pronounced or extended detachments from the NL. To determine the expression level relative to endogenous genes, we performed RNA-seq. Transcriptional activity was readily detectable for LAD integrations, although their median expression was about 4-fold lower compared to the iLAD integrations (**Figure 8d**). Taking this difference into account, and after correction for the copy number in each clone, the median expression level of the integrated transgenes inside LADs still ranks approximately in the upper 97^th^-98^th^ percentile of all active endogenous genes (**Figure 8e**). Thus, even within LADs most of the integrated transgenes are expressed at very high levels. These expression levels can be sufficient to reduce NL interactions, but only moderately and locally.

## DISCUSSION

Evidence that components of the transcription machinery can affect the spatial organization of the genome is accumulating, but the underlying processes are still poorly understood (van Steensel and Furlong, 2019; Vermunt et al., 2019). The data presented here consistently show that activation of genes in LADs leads to detachment from the NL, and conversely that inactivation can lead to increased NL contacts. Moreover, the results point to a remarkable flexibility of the chromatin fiber, allowing for the repositioning of individual genes without much effect on flanking DNA.

Several of our results point to a role for transcription elongation in counteracting NL interactions. First, activity-induced detachment from the NL generally extends across the entire activated transcription unit, from the activated promoter until the 3’ end of the gene. We observed this for a wide range of gene sizes. A role for elongation is also strongly supported by premature termination of the active *Cobl* gene by insertion of a PAS, which primarily caused an increase of NL interactions downstream but not upstream of the new termination site. These results are consistent with a study of the *ThymoD* non-coding RNA gene in mouse T cell progenitors, where insertion of a PAS prevented detachment from the NL as observed by FISH (Isoda et al., 2017). Conversely, read-through transcription into heterochromatin, elicited by influenza virus NS1 protein, was found to cause relocation from the heterochromatic compartment “B” to the euchromatic compartment “A” (Heinz et al., 2018), which largely correspond to LADs and iLADs, respectively (van Steensel and Belmont, 2017). How transcription elongation may prevent NL interactions remains to be elucidated. It could be a physical effect, for example when a transcribed gene is tethered to a structure in the nuclear interior. It may also be a biochemical effect, such as the removal of particular NL-interacting chromatin proteins by the elongating RNA polymerase complex.

Earlier work found that VP16-induced movement of a LacO repeat towards the nuclear interior could not be blocked by elongation inhibitors (Chuang et al., 2006). This is not necessarily contradictory to our evidence that supports a role for elongation; it is possible that a transcription activator like VP16 also promotes detachment from the NL independently of transcription elongation. In support of such an additional mechanism, some of the genes that we studied (e.g. **Figures 1a, 2a, 6a**) showed the strongest loss of NL interactions near their 5’ end. Similarly, a class of naturally active genes inside LADs exhibits more prominent detachment of their TSS compared to the downstream transcription units (Leemans et al., 2019; Luperchio et al., 2017; Wu and Yao, 2017). Furthermore, global tethering of VP64 across all LADs caused virtually no changes in transcription yet triggered loosening of LAD-NL interactions (Kind et al., 2013), underscoring that VP16 can counteract NL interactions without activating transcription.

In two earlier reports, relocation from the NL to the nuclear interior was also achieved by tethering of an artifical peptide that induces chromatin decondensation without detectable recruitment of RNA Polymerase II (Chuang et al., 2006; Therizols et al., 2014). It is not understood how this peptide (which is not derived from a naturally occurring protein) exerts this effect, but it suggests that a transcription-independent mechanism of relocation exists in addition to transcription-linked mechanisms. Recent evidence suggests that active chromatin marks such as H3K27ac deposited by p300 may counteract NL interactions (Cabianca et al., 2019).

In most cases, the transcriptionally inactive regions adjacent to our activated genes remain relatively unaffected in their NL contacts. Conversely, inhibition of transcription (either by deleting promoters or by insertion of a PAS) leads to increased NL interactions in a very local manner. The latter results were obtained in genomic regions that are facultative LADs, i.e., they may have an intrinsic ability to interact with the NL in the absence of transcription. Genes in constitutive iLADs may lack this ability, either due to spatial constraints or because they lack certain sequence features or chromatin characteristics. It will be of interest to further dissect the molecular mechanisms that underlie the apparent competition between forces that tether chromatin to the NL and forces (such as transcription elongation) that counteract these interactions.

These and previously reported data (de Leeuw et al., 2018; Gonzalez-Sandoval and Gasser, 2016; Kim et al., 2019; Lochs et al., 2019; van Steensel and Belmont, 2017) together suggest a balancing act between transcription and LADs: for many genes, LADs pose a repressive environment (Leemans et al., 2019). This, however, may be overcome by strong transcription activators. Once transcription is active, it causes detachment of the gene from the NL. Possibly this helps to reinforce the active state. It was previously found that a transiently activated gene can remain detached from the NL for several days after the activation signal has subsided (Therizols et al., 2014).

We observed changes from late to early replication timing for all five upregulated genes that we assayed by Repli-Seq. The regions that exhibited shifts in replication timing roughly match in size with the usual span of replication domains observed in vivo (Hiratani et al., 2008; Pope et al., 2014), suggesting that there may be a fundamental minimal size of such domains. Interestingly, the overlap with changes in NL interactions, while substantial, was imperfect. The shifts in replication timing tended to involve a larger region than the shift in NL interactions, and were centered around the targeted promoters rather than the entire transcription unit. Together, these results suggest that both NL interactions and replication timing can be modulated by the transcription machinery, but elongation appears to to play a more prominent role in counteracting NL interactions, while a signal emanating from activated promoters may evoke a change in replication timing. These distinct but closely linked mechanisms may explain why LADs and late-replicating domains overlap strongly but imperfectly.

## Acknowledgments

We thank the NKI Genomics, Flow Cytometry and RHPC core facilities, as well as Tom Rieuwerts and Ludo Pagie for technical assistance. We thank Andrew Belmont, Jian Ma and other members of the 4DN Center for Nuclear Cytomics for helpful discussions. Supported by NIH Common Fund “4D Nucleome” Program grant U54DK107965 (BvS and DMG) and European Research Council Advanced Grant 694466 (BvS). The Oncode Institute is partly supported by KWF Dutch Cancer Society.

## Author contributions

LB: Conceived and designed study, conducted majority of experiments, initial data analysis, wrote manuscript. PAZ: Performed Repli-seq experiments and initial Repli-seq data analysis. DPH: Performed experiments, and data analysis. TvS: Processing of DamID data. CL: Performed data analysis. JS: *Morc1* locus deletions and initial data analysis. DMG: Supervised Repli-seq experiments, generation of *Morc1* locus deletions and initial data analysis. BvS: Designed study, performed coding and data analysis, wrote manuscript, supervised project.

## MATERIALS AND METHODS

### Cell culture

The RPE-1 cell line stably expressing SunTag-CRISPRa (Tame et al., 2017) was kindly provided by the R. Medema lab (Netherlands Cancer Institute, Amsterdam) and cultured in DMEM-F12 supplemented with 10% FCS. F121-9 mES cells were kindly provided by J. Gribnau (Erasmus Medical Center, Rotterdam, the Netherlands) and cultured in feeder-free 2i medium according to the 4DNucleome protocol (https://data.4dnucleome.org/protocols/cb03c0c6-4ba6-4bbe-9210-c430ee4fdb2c/).

### TALE-VP64 experiments

TALE-VP64 constructs (Therizols et al., 2014) with Puro resistance marker were kindly provided by Pierre Therizols.

F121-9 cells were transfected with TALE-VP64 constructs targeting Nrp1 or Sox2 by electroporation using Lonza Mouse Embryonic Stem Cell Nucleofector™ Kit (VPH-1001) according to the manufacturer’s instructions. Cells were selected with Puromycin (1 µg/µl) for one week to obtain stable polyclonal cell pools.

### CRISPRa experiments

sgRNAs were cloned into Lentiguide-Puro vector (Addgene #52963) using restriction enzyme BsmBI, and Lentivirus was prepared. RPE-1 cells stably expressing SunTag-CRISPRa were infected with LentiGuide virus and selected with 10 µg/µl Puromycin for one week to obtain stable polyclonal cell pools.

### PAS integration

A PAS was inserted by in-frame integration of a blasticidin resistance (BlastR) cassette followed by the PAS into the *Cobl* gene. For this purpose, sgRNA sequence AGTCATCTGTGCGAAGTGTG was cloned into Blast-TIA vector (Lackner et al., 2015) (kindly supplied by the Brummelkamp lab) via BbsI restriction digestion. Cells were transfected with the resulting Blast-TIA vector co-transfected with Cas9 expression vector pX330 (Addgene #42230) by nucleofection and subjected to selection by culturing in the presence of 10 µg/µl Blasticidin for one week. Clones were picked and screened for correct integration of the BlastR cassette by PCR with primers lb877 and lb982. Heterozygosity of the integration was confirmed by PCR using primers lb982 and lb991, and the *129Sv* allele was identified as the targeted allele by PCR using primers lb877 and lb1010, followed by Sanger sequencing with the same primers.

### Repli-seq

Repli-seq was performed as described (Marchal et al., 2018). Sequencng was done on a NovaSeq 6000 system (Illumina), 50 bp read length.

### RNA-seq

Strand-specific mRNA-seq was performed by selection of polyadenylated transcripts, reverse transcription and sequencing on an Illumina HiSeq2500 sequencer. Reads were aligned to hg19 or mm10 using TopHat version 2.1, with Ensembl genome build 75. For *Cobl* PAS integration experiments in the F1 hybrid ES cells, mRNA-seq reads were aligned to mm10 using STAR (Dobin et al., 2013).

### RT-qPCR

Cells were collected in TRIsure and total RNA was extracted using PureLink RNA Mini Kit (Thermo Fisher Scientific) according to the manufacturer’s instructions. RNA was reverse-transcribed using Tetro Reverse Transcriptase (Bioline) with Oligo(dT)20 primers (Thermo Fisher Scientific) according to the manufacturer’s instructions. qPCR was performed using SensiFast no-ROX mix (Bioline) in a 10 µl reaction. Primers are listed in Table S3.

### Generation and mapping of random integrations

The hPGK-EGFP cassette was derived from plasmid TRIP vector pPTK-Gal4-mPGK-Puro-IRES-eGFP-sNRP-pA (Akhtar et al., 2014) by replacing mPGK-Puro-IRES with the human PGK promoter using restriction enzyme cloning with SalI and NcoI. Generation of a barcoded plasmid pool and integration into F121-9 mES cells was performed as described (Akhtar et al., 2014). Clones with high EGFP expression were sorted by FACS and screened for high integration copy number by qPCR with EGFP-specific primers EB66 and EB38, using *Lbr*-specific primers (JOYC231 and JOYC232) for normalization.

Mapping of all integrations without linking to barcodes was done by Tagmentation as described (Stern, 2017) with minor modifications: Before PCR for Tn5 adapters, linear amplification of PiggyBac integrations was performed using primers lb565 or lb566 for mapping in revese or forward orientation respectively. Linear amplification was performed using 0.5 U Phusion Polymerase (Bioline) in a 20µl reaction with Phusion GC-rich buffer, 1mM dNTPs, 50nM primer. Reaction was incubated at 98°C for 30 sec, then 45 cycles of 98°C for 8 sec, 60°C for 5 sec and 72°C for 30 sec followed by a final step at 72°C for 20 sec. For PCR amplification, PiggyBac-specific primers lb565 or 566 were used for mapping in reverse or forward orientation respectively.

To process the tagmentation mapping reads, the Tn5 adaptor sequence and PiggyBac primer sequence at the ends of the paired-end reads were removed using an adaptation of c*utadapt v1.11*. The genomic part of the sequence was mapped to strain-specific versions of GRCm38 release 68 from Ensembl using *bowtie v2.3.4.1* with mapping set to “*very-sensitive*”. To create these strain-specific genomes, SNP information was downloaded from the Mouse Genomes Project (Keane et al., 2011) (http://www.sanger.ac.uk/science/data/mouse-genomes-project) as VCF files “CAST_EiJ.mgp.v5.snps.dbSNP142.vcf.gz” and “129S1_SvImJ.mgp.v5.snps.dbSNP142.vcf.gz” (version 1 May 2015). *Bcftools* was used to incorporate all SNPs into the GRCm38 reference genome. After mapping to strain-specific genomes, bam files were compared and for each read, the alignment with the highest alignment score (AS) was used. When the AS was identical, a random choice was made. Read-pairs aligning in opposite orientation and less than 1500 bp apart were converted to genomic regions using the *bamToBed* from *bedtools* and *awk,* covering both reads as well as the region in between. *Genomecov* from *bedtools* and *awk* was used to combine regions and calculate coverage. Integration sites were called by combining regions from PCRs from both transposon arms using *closest* from *bedtools* and *awk*. Regions on opposite strands that were at most 5 bp apart were regarded to represent an integration. Next, the allele of the integration was determined by using *mpileup* from *Samtools* (Li et al., 2009) *v1.5* with a maximum depth of 50 to count the number of mismatch positions over the complete region compared to both of the strain-specific GRCm38 modifications. Each position with the allele of the strain-specific genome occurring in a ratio less than 0.5 was considered a mismatch position. The allele with the lowest number of mismatch positions was then considered the allele of integration. In case of equal number of mismatch positions, the integration allele was classified as ambiguous.

In addition, allele-specific mapping of the integrations and linking to their barcodes was performed by inverse PCR as described (Akhtar et al., 2014), except that the mapping of reads was confined to regions that were initially found by tagmentation mapping. This identified a subset of 63 integrations that could be linked to a barcode. In total, we identified ## mapped integrations for clone CM1407 and ## for clone CM1420, of which ## and ## were linked to a barcode, respectively.

### Expression analysis of ES cell clones with random integrations

Clones CM1407 and CM1420 were subjected to mRNA-seq as above. For comparison between eGFP and endogenous mRNA expression, a fasta entry for eGFP was added to the mouse genome version mm10 chromosomes 1-19, X, Y and M without alternative contigs. The annotation of eGFP transcript was also added to gencode version M18. STAR (Dobin et al., 2013) version 2.6.0c was used to align the cDNA reads to this modified reference genome and for each transcript reads were counted. DESeq2 (Love et al., 2014) was used to calculate Fragments Per Kilobase Million (FPKM) values for each gene, including eGFP. In addition, barcode-specific expression was determined using the standard TRIP protocol (Akhtar et al., 2014). Based on the genomic DNA barcode counts, the number of integrations per clone was calculated. This was done by first selecting all genuine barcodes sequenced > 500 times in the sample. To discriminate between genuine barcodes and sequencing errors of these barcodes, *starcode* (Zorita et al., 2015) was used. Since a single barcode can be integrated multiple times, these barcode counts were divided by the median barcode count. This value was then rounded to the nearest integer, estimating the number of integrations per barcode. For clone CM1420 this resulted in an estimation of 51 barcodes with single integrations and 2 barcodes with 2 integration sites. For the clone CM1417, 44 single barcode integrations were estimated and 3 barcodes were estimated to be integrated at 2 locations. Finally, the eGFP FPKM was scaled by the number of integrations for each clone (55 and 50 respectively) in order to determine the average eGFP expression per integrated reporter.

LAD coordinates in mouse ES cells were obtained from ref (Peric-Hupkes et al., 2010) and adjusted to mm10 using the LiftOver tool (Hinrichs et al., 2006). To estimate FPKM’s for integrations in LADs and iLADs separately, barcodes were used that had a unique location according to a combination of iPCR and tagmentation mapping. In total, 50 barcodes could be confidently linked to iLAD locations and 13 in LADs. To determine LAD and iLAD-specific expression levels, the average eGFP FPKM per integration was scaled by the median LAD and iLAD expression. Finally, the percentile of eGFP FPKM relative to endogenous active genes was calculated by counting the number of genes with higher FPKM than the eGFP estimation, divided by the total number of active genes (defined as genes with FPKM > 0; n=26128 for clone CM1420 and n=27688 for clone CM1417).

### DamID-seq

DamID-seq was performed as described (Brueckner et al., 2016) with minor modifications. Dam fused to human LMNB1 protein (Dam-LMNB1) or unfused Dam were expressed in cells by lentiviral transduction (Vogel et al., 2007). Three days after infection, cells were collected for genomic DNA (gDNA) isolation. gDNA was pre-treated with SAP (10 U, New England Biolabs #M0371S) in CutSmart buffer in a total volume of 10 µl at 37 °C for 1 h, followed by heat-inactivation at 65°C for 20 min to suppress signal from apoptotic fragments. This gDNA was then digested with DpnI (10 U, New England Biolabs #R0176L) in CutSmart buffer in a total volume of 10 µl at 37 °C for 8 h followed by heat inactivation at 80 °C for 20 min. Fragments were ligated to 12.5 pmol DamID adapters using T4 ligase (2.5 U, New England Biolabs ##) in T4 ligase buffer in a total volume of 20 µl incubated at 16 °C for 16 h. The reaction was heat-inactivated for 10 min at 65 °C. Products were then digested with DpnII to destroy partially methylated fragments. DpnII buffer and DpnII (10 U, New England Biolabs #R0543L) were added in a total volume of 50 µl and incubated at 37 °C for 1 h. Next, 8 µl of DpnII-digested products was amplified by PCR with MyTaq Red Mix (Bioline #BIO-25044) and 1.25 µM primers Adr-PCR-Rand1 in a total volume of 40 µl. PCR settings were 8 min at 72 °C (1×) followed by 20 s at 94 °C, 30 s at 58 °C, 20 s at 72 °C (24× for Dam, 28x for Dam-LMNB1 samples) and 2 min at 72 °C (1×). Remaining steps were performed as previously described. Samples were sequenced on an Illumina HiSeq2500.

### Processing of RPE-1 and ES cell DamID data

First, the constant DamID adapter was trimmed from the 65 bp single-end reads using cutadapt (Martin, 2011) version 1.11 and custom scripts. The remaining sequence starting with GATC was mapped to hg19 with bowtie2 (Langmead and Salzberg, 2012) version 2.2.6. Uniquely mapped reads (filtered for bowtie’s XS-tag) were then assigned to individual gDNA sequences between two GATC motifs (referred to as GATC fragments), which are the units of further data processing and analysis because Dam only methylates GATC motifs. Across all RPE-1 experiments, the median number of reads mapped to GATC fragments was ## (Dam-LaminB1) and ## reads (Dam-only). Further processing and analysis was done in R (R Core Team., 2017) versions 3.4 - 3.6 using Bioconductor (Huber et al., 2015), in particular the packages GenomicRanges (Lawrence et al., 2013) and Sushi (Phanstiel et al., 2014).

Replicate experiments were combined by summing the reads for each GATC fragment. Hence, experiments with more reads were weighed proportionally stronger than experiments with fewer reads. Extremely high read counts of individual GATC fragments (>100 times the genome-wide average) were assumed to be due to PCR artefacts; these read counts were replaced with the genome-wide average read count. Next, smoothing was applied by summing read counts over a running window of 201 consecutive GATC fragments. A pseudocount of 30 was added to each window. The ratio Dam-LaminB1 / Dam-only was calculated for each window and log_2_ transformed. Finally, the log_2_ ratios were normalized by subtracting the genome-wide average log_2_ ratio.

When comparing experimental and control DamID logratios in genome-wide scatterplots, we noticed modest systematic biases and skews (visible as point clouds that were somewhat banana-shaped rather than cigar-shaped) or differences in the dynamic ranges of the DamID values that are likely to be of technical nature (under the assumption that CRISRPa activation of a single gene is unlikely to cause a genome-wide systematic effect). We estimated such skews empirically by applying a lowess fit (span = 0.5) to the experimental ∼ control comparison of a random selection of 50,000 GATC fragments, and then used this fit to correct the genome-wide comparison. This effectively removed genome-wide biases, thereby enhancing the sensitivity to detect local changes in NL contacts around the targeted genes.

### Processing of F1 hybrid mouse ES cell DamID data

For DamID on F1 hybrid *129*/*Cast* mouse ES cells, 150 or 200 nt single reads were trimmed to remove the DamID adaptor sequence, and then strain-specifically mapped to mm10 with WASP (van de Geijn et al., 2015) using bowtie2 and VCF files from the mouse genomes project (Keane et al., 2011) (https://www.sanger.ac.uk/science/data/mouse-genomes-project; version 5). Data was further processed as described for RPE-1 cells, except that a smoothing window size of 301 instead of 201 GATC fragments was applied.

### Domainograms

The domainograms in this study are related to those reported previously (de Wit et al., 2008; Tolhuis et al., 2011), but do not show estimated P-values, which are not easily calculated for our experimental design. Rather, for a given window size, they show the ranking of changes in DamID log-ratios (experimental minus control) relative to windows of the same size genome-wide. Briefly, in a window of *w* neighboring GATC fragments, the difference in mean DamID log-ratio is calculated between the experimental and control samples. This is done for all possible windows of size *w* genome-wide. Next, windows in which both experimental and control sample showed only baseline DamID signals (i.e., both log-ratios are in the respective lower 0.3 quantiles genome-wide) are discarded. This is done because baseline fluctuations can appear strong on a logarithmic scale but are generally of minor amplitude on a linear scale, and therefore unlikely to be of biological relevance. The remaining windows are ranked by their log-ratio differences; ranks <5% or >95% are visualized by blue or red color scales, respectively. This is repeated for 28 different window sizes *w* that are logarithmically ranging from 67 and 2917 GATC fragments, i.e. from ∼30kb to ∼1Mb.

### Data analysis of Repli-seq samples

Repi-seq reads from early and late replicating fractions were mapped and processed in the same way as DamID reads, using the same smoothing window size. Instead of Dam-LaminB1/Dam-only, the ratio late/early replication was calculated.

## Data availability

Sequencing reads and processed data of DamID, Repli-seq and RNA-seq experiments are available from the Gene Expression Omnibus (https://www.ncbi.nlm.nih.gov/geo/), accession GSE133275.

## SUPPLEMENTARY FIGURE LEGENDS

**Figure S1. Expression of genes in RPE-1 cells targeted by CRISPRa.** Expression levels were determined by RT-qPCR. Average of three technical replicates, error bar indicates standard deviation. fc/GAPDH: expression level normalized to *GAPDH* gene. ctrl: untransfected control.

**Figure S2. Expression changes of CRISPRa-targeted genes and neighboring genes in RPE-1 cells.** Top panels: domainograms showing changes in NL interactions around genes activated by CRISPRa, similar to **Figure 2**. Middle panels: log_2_ changes in gene expression as determined by mRNA-seq, compared to untransfected control cells (average of two independent replicate experiments each). Data for *ADAM22*, *ABCB1* and *PTN* are identical to those in **Figures 2c** and **4**. Bottom panels: gene annotation track. Each CRISPRa-targeted gene is highlighted in green; targeted promoters are marked by vertical green dotted line.

**Figure S3**: **Changes in NL interactions and replication timing in the CCSER1/GRID2 locus**. Effects of CRISPRa activation of *CCSER1* **(a)**, *GRID2* **(b)** or both genes **(c)** in RPE-1 cells. Top panels show DamID data similar to **Figure 2a-b**. Middle panels show maps of replication timing at the same resolution and in the same plotting style as for DamID, except that different colors are used as indicated in **(d)**. Bottom panels show gene annotation; activated gene(s) highlighted in green. DamID data are the same as in **Figures 2c** and **6a**.

**Figure S4: Changes in NL interactions of genes upregulated by CRISPRa**. DamID data obtained after CRISPRa in RPE-1 cells of *SLC35F3* **(a)**, *TRAM1L1* **(b)**, *ZNF804B* **(c)**. Domainograms are the same as in **Figure 2c**, but additionallly show increased NL contacts in red.

**Figure S5: Expression and NL interactions of** *ABCB4*, *ABCB1*, *RUNDC3B* **and** *ADAM22* **after combined activation**. **(a)** CRISPRa was done using four sgRNAs simultaneously, one for each promoter. Expression was quantified by mRNA-seq. Results from two independent replicates are shown. ND, not detectable. **(b)** Visualization of log_2_ changes in gene expression (middle panel; average of the same two experiments as in (a)), together with DamID domainogram (top panel) and gene annotation track (bottom panel). CRISPRa-targeted genes are highlighted in green, and the sgRNA target locations are marked by vertical green dotted lines. **(c-e)** DamID profiles after activation of *ADAM22* (**c**), *ABCB1* (**d**), or *ABCB4*, *ABCB1*, ADAM22 and *RUNDC3B* simultaneously (**e**). Data visualization as in **Figure 2a-b**.

**Figure S6: Effect of PAS integration on** *Cobl* **gene transcription.** Ratio of allele-specific mRNA-seq read counts of *129Sv* and *Cast* SNP variants located in the *Cobl* gene. Two SNPs are located upstream of the PAS integration site; all other SNPs are downstream. Note that the 5’ to 3’ orientation of the gene is from right to left. In a control clone without the PAS, the *Sv129* allele is consistently expressed ∼2-fold higher than the *Cast* allele. In the clone with the PAS integration in the *129Sv* allele, expression of the *129Sv* variants downstream of the PAS is strongly reduced. The increased level of the *129Sv* variants upstream may be due to greater stability of the mRNA due to its altered 3’ end. Data are average of two independent biological replicates.

## Supplementary Tables

**Table S1:**
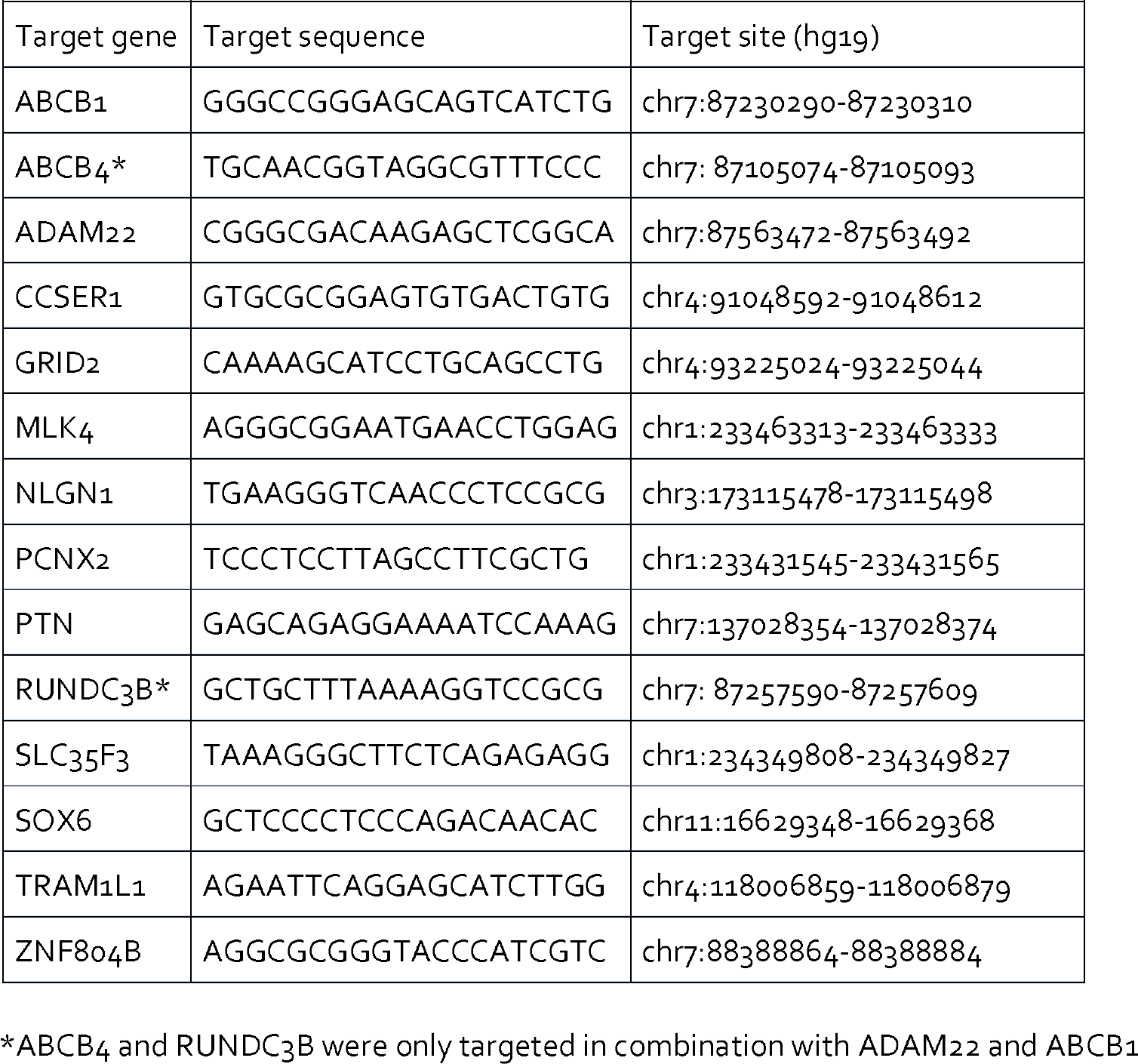
sgRNA sequences CRISPRa

**Table S2:**
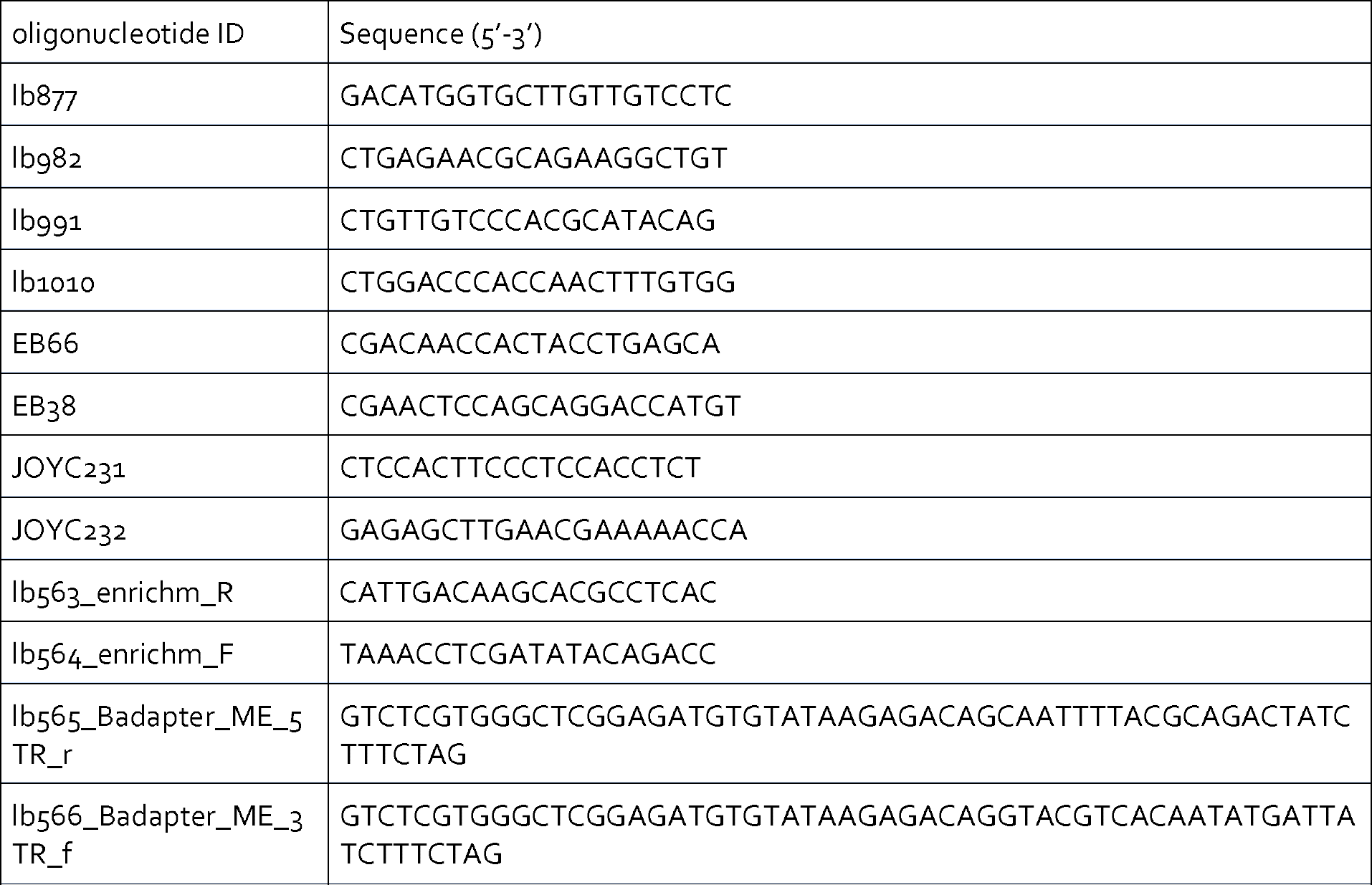
Miscellaneous oligonucleotides used.

**Table S3:**
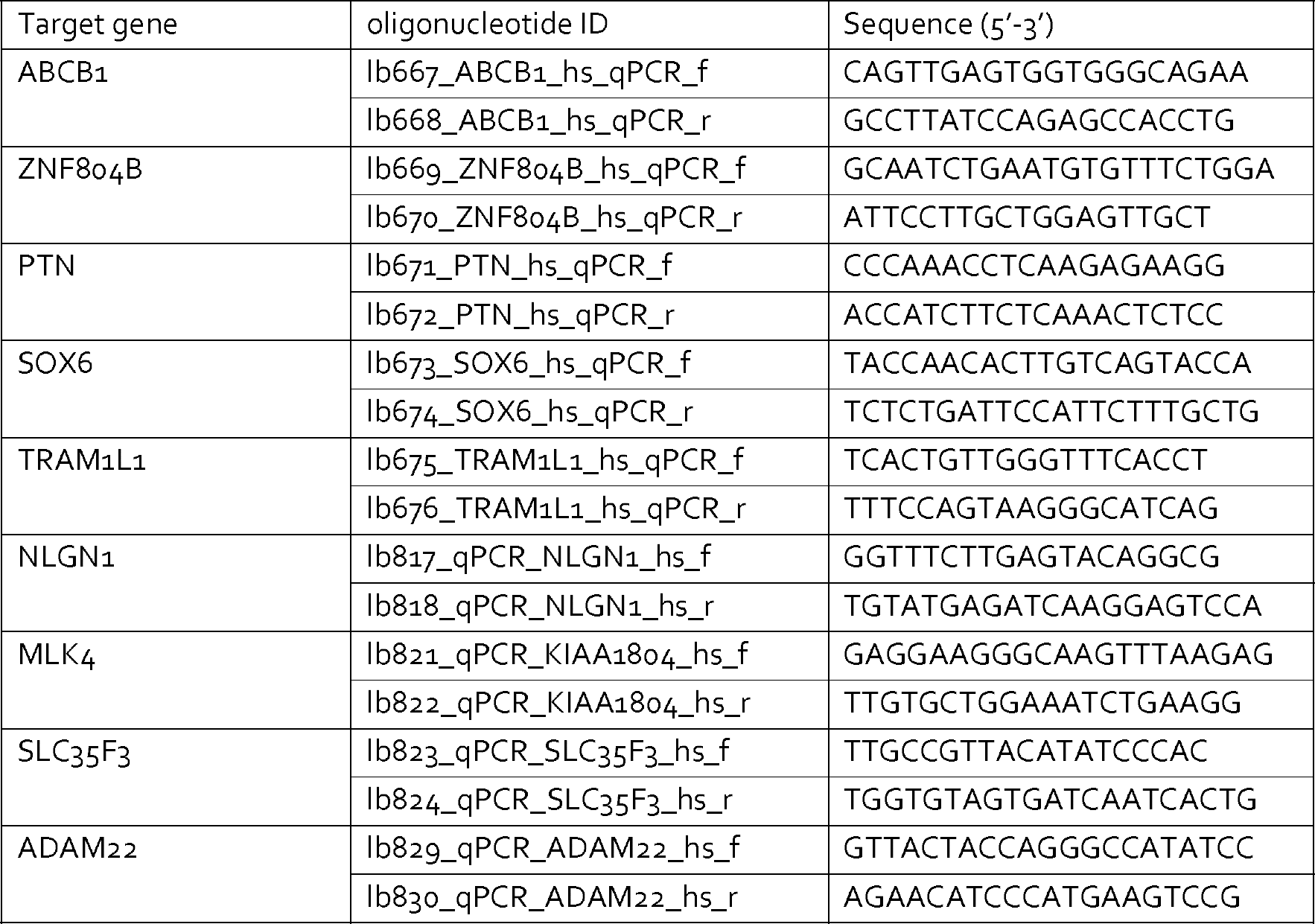
Oligonucleotides for RT-qPCR.

