## Supplementary figures and images for "Local rewiring of genome - nuclear lamina interactions by transcription"

### Supplemental Figures S1-S6

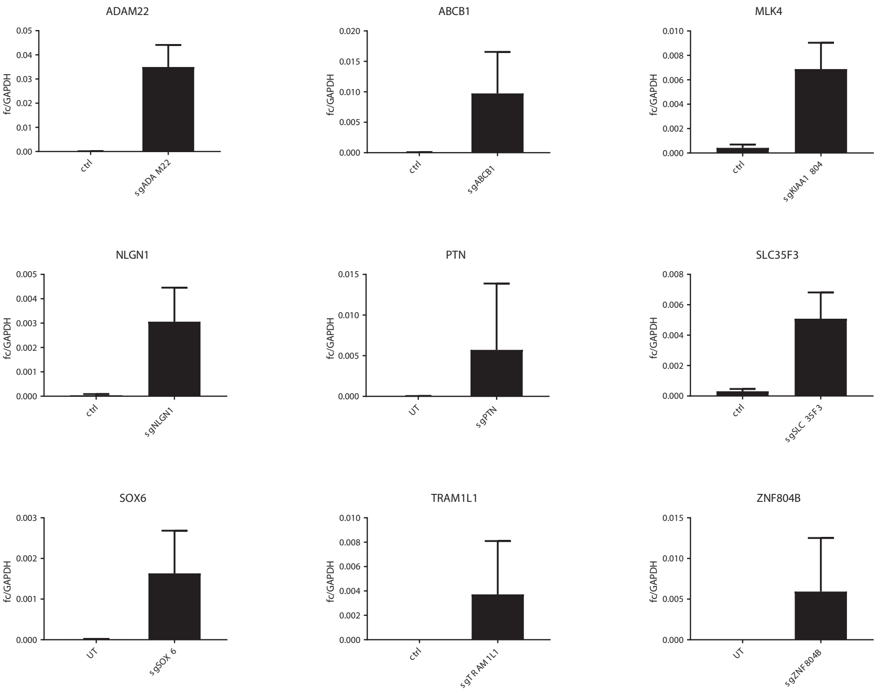

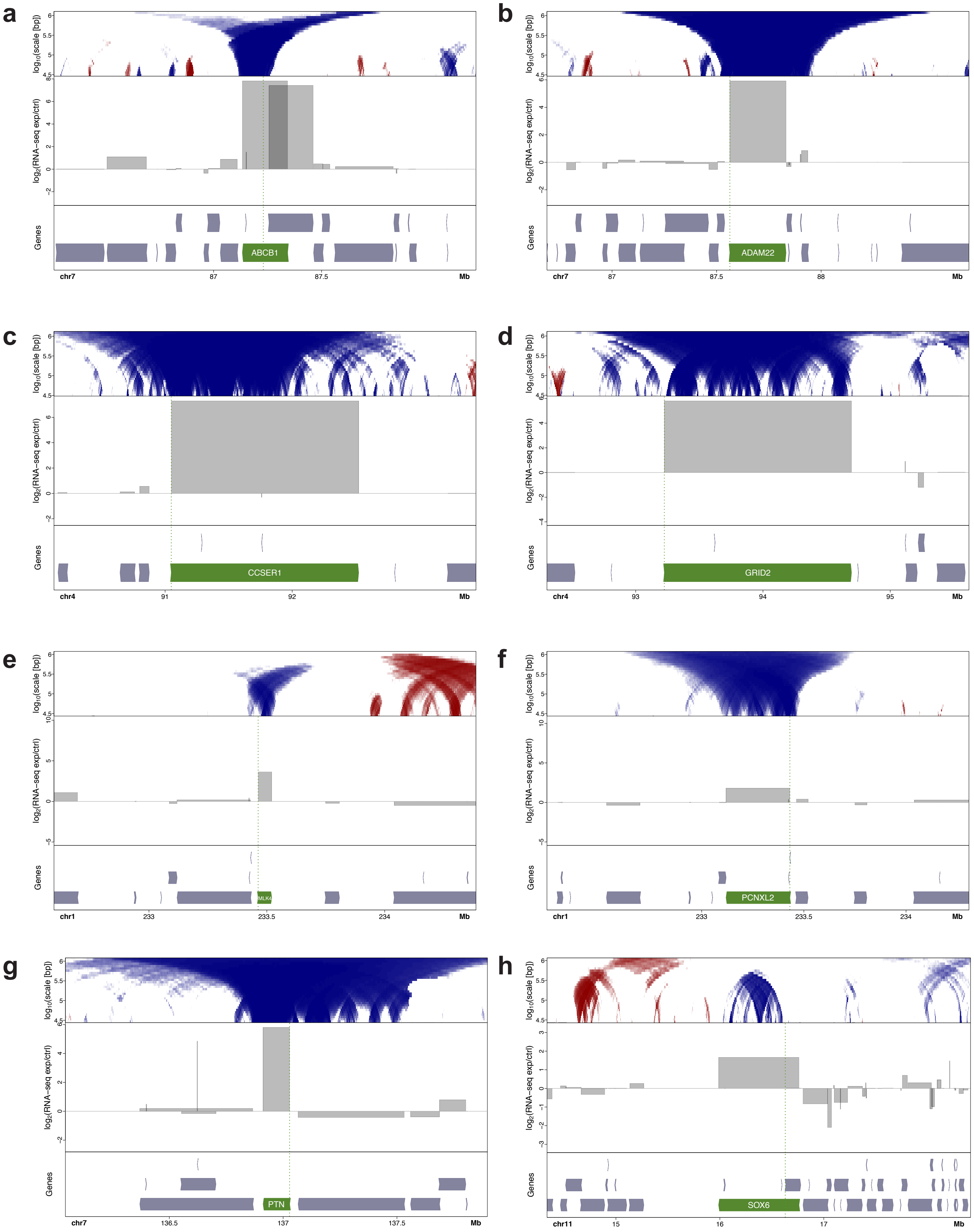

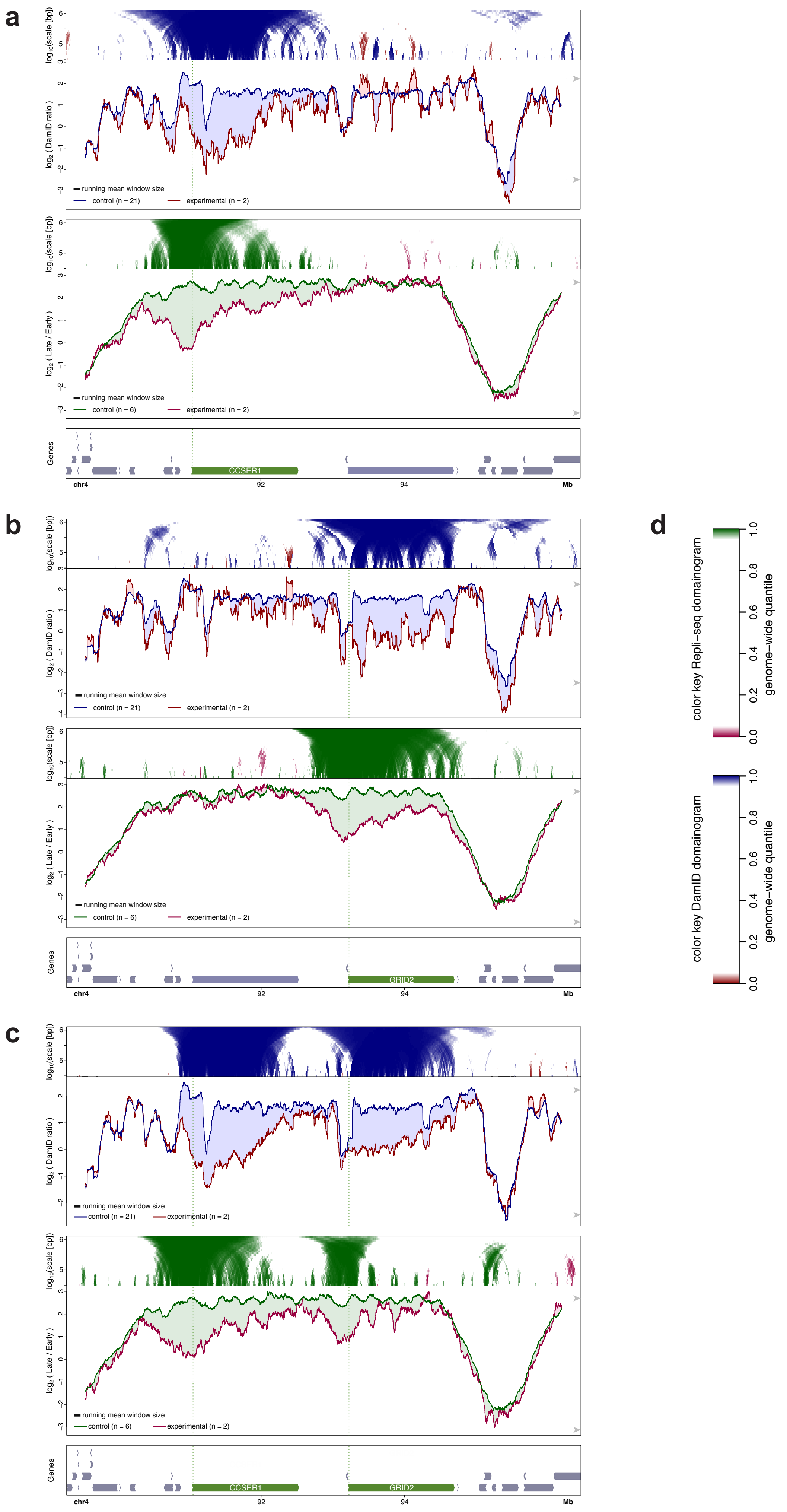

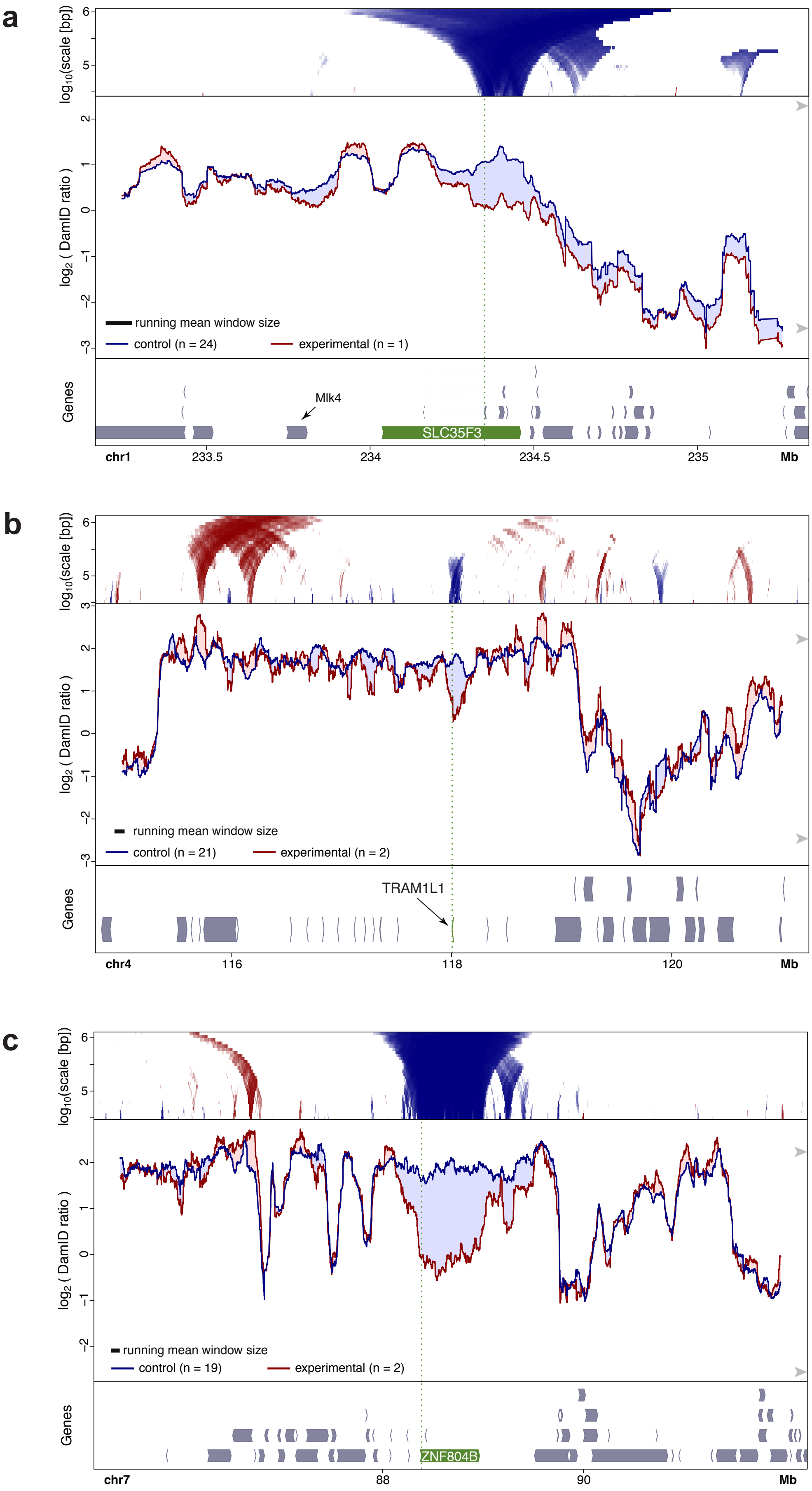

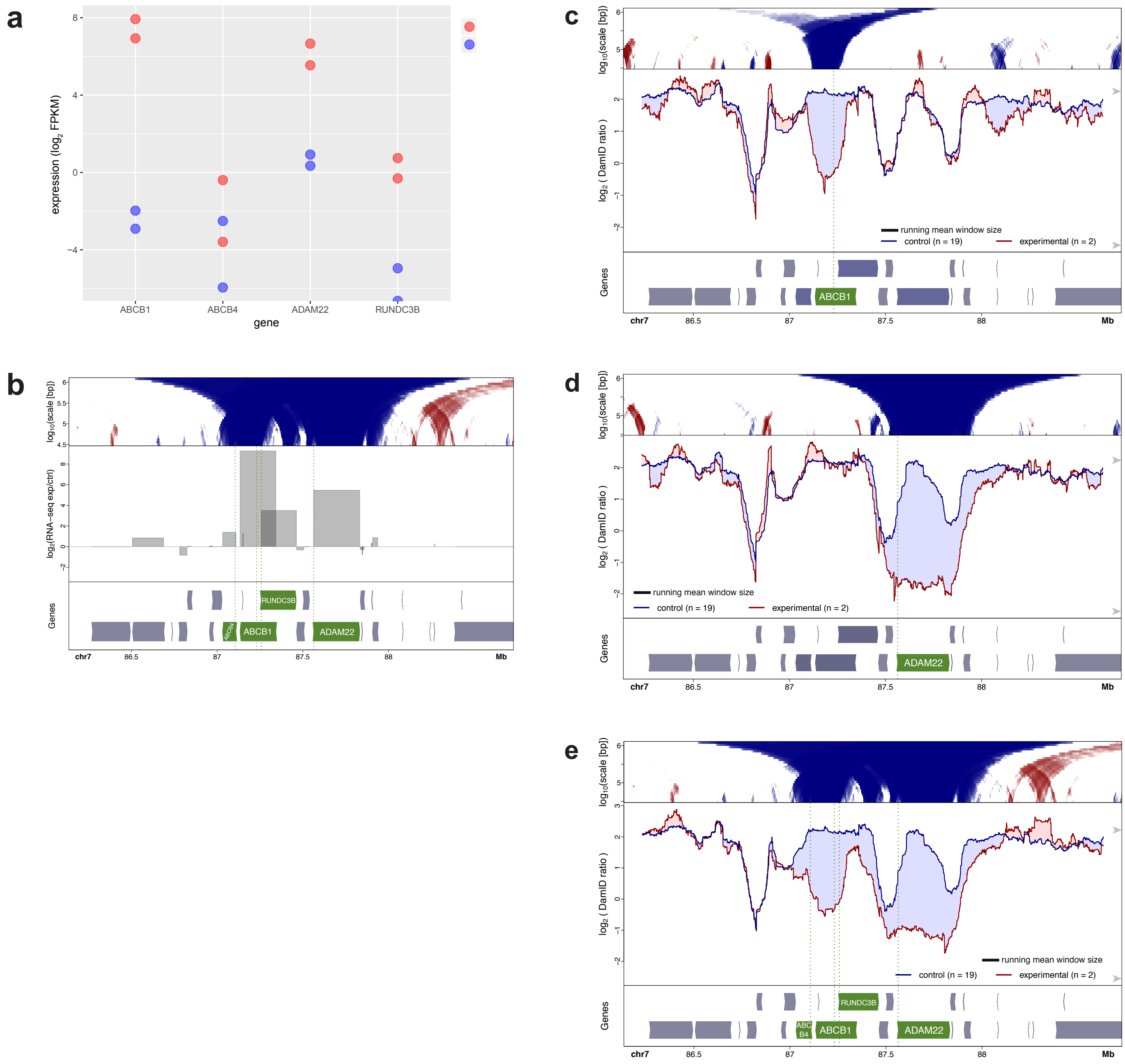

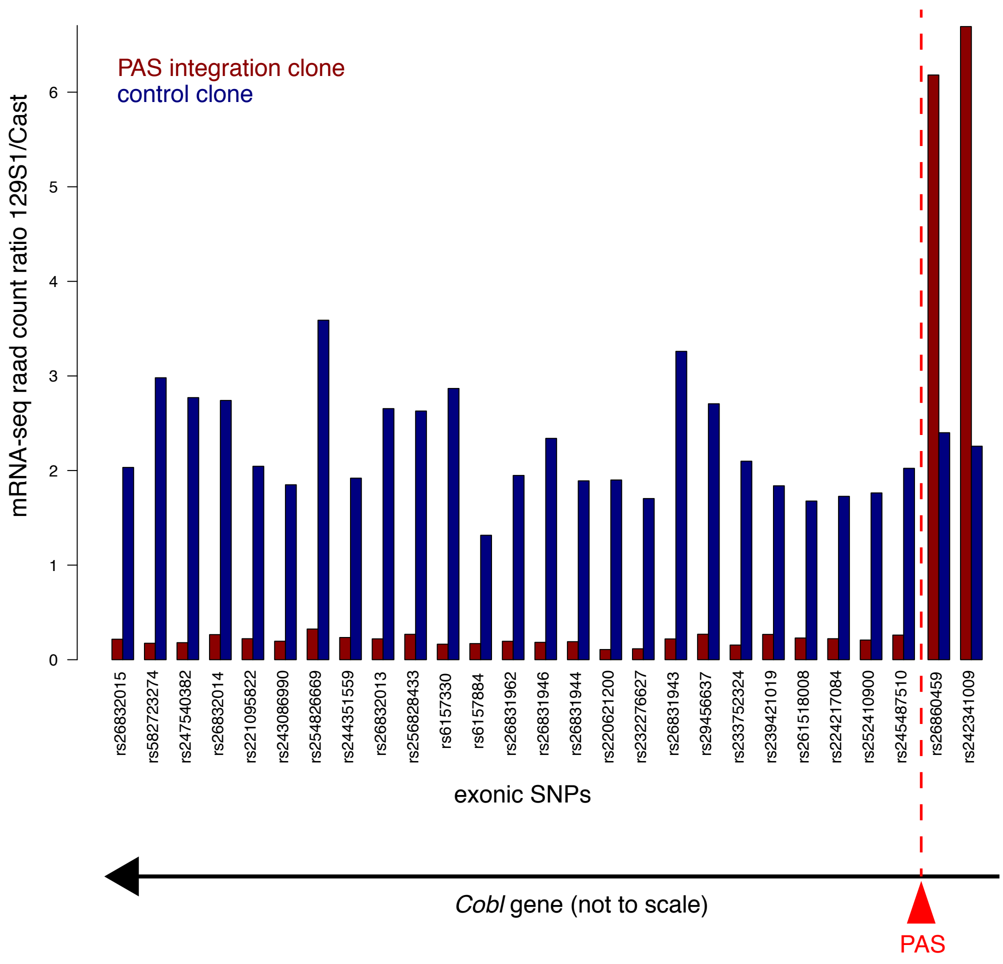
